# Temperature impacts on fish physiology and resource abundance lead to faster growth but smaller fish sizes and yields under warming

**DOI:** 10.1101/2021.10.04.463018

**Authors:** Max Lindmark, Asta Audzijonyte, Julia Blanchard, Anna Gårdmark

**Author notes:** Author to whom correspondence should be addressed. Current address: Max Lindmark, Swedish University of Agricultural Sciences, Department of Aquatic Resources, Institute of Marine Research, Turistgatan 5, 453 30 Lysekil, Sweden.

## Abstract

Resolving the combined effect of climate warming and exploitation in a food web context is key for predicting future biomass production, size-structure, and potential yields of marine fishes. Previous studies based on mechanistic size-based food web models have found that bottom-up processes are important drivers of size-structure and fisheries yield in changing climates. However, we know less about the joint effects of ‘bottom-up’ and physiological effects of temperature; how do temperature effects propagate from individual-level physiology through food webs and alter the size-structure of exploited species in a community? Here we assess how a species-resolved size-based food web is affected by warming through both these pathways, and by exploitation. We parameterize a dynamic size spectrum food web model inspired by the offshore Baltic Sea food web, and investigate how individual growth rates, size-structure, relative abundances of species and yields are affected by warming. The magnitude of warming is based on projections by the regional coupled model system RCA4-NEMO and the RCP 8.5 emission scenario, and we evaluate different scenarios of temperature dependence on fish physiology and resource productivity. When accounting for temperature-effects on physiology in addition to on basal productivity, projected size-at-age in 2050 increases on average for all fish species, mainly for young fish, compared to scenarios without warming. In contrast, size-at-age decreases when temperature affects resource dynamics only, and the decline is largest for young fish. Faster growth rates due to warming, however, do not always translate to larger yields, as lower resource carrying capacities with increasing temperature tend to result in declines in the abundance of larger fish and hence spawning stock biomass. These results suggest that to understand how global warming affects the size structure of fish communities, both direct metabolic effects and indirect effects of temperature via basal resources must be accounted for.

## Introduction

Climate change affects aquatic food webs directly by affecting species’ distribution (Pinsky *et al*. 2013), abundance (McCauley *et al*. 2015), body size (Daufresne *et al*. 2009; Baudron *et al*. 2014), and ecosystem function (Pontavice *et al*. 2019). Global retrospective analysis of warming and fish population dynamics has revealed that the maximum sustainable yield of scientifically assessed fish populations across ecoregions has already declined by 4.1% on average between 1930 and 2010 due to climate change (Free *et al*. 2019). These results are also matched in magnitude and direction by projections from an ensemble of mechanistic ecosystem models, which predict ∼5% decline in animal biomass for every 1 °C of warming, especially at higher trophic levels (Lotze *et al*. 2019). Across a range of process-based ecosystem models, declines in productivity of fish stocks and abundance of large fish have been linked to changes in primary production or zooplankton abundance (Blanchard *et al*. 2012; Woodworth-Jefcoats *et al*. 2013, 2015; Barange *et al*. 2014; Lotze *et al*. 2019; Heneghan *et al*. 2021; Tittensor *et al*. 2021). However, even in areas where warming is predicted to have positive effects on primary production, fish productivity does not appear to increase (Free *et al*. 2019). This suggests that fish population dynamics might be strongly influenced by other factors, such as temperature-driven changes in recruitment, mortality or somatic growth (Free *et al*. 2019), yet the driving mechanisms remain poorly understood.

Global warming is also predicted to cause reductions in the adult body size of organisms, and this is often referred to as the third universal response to warming (Daufresne *et al*. 2009; Sheridan & Bickford 2011; Forster *et al*. 2012). It is often attributed to the temperature-size rule (TSR) is observed in a wide range of ectotherms (Forster *et al*. 2012). This is an intraspecific rule stating that individuals reared at warmer temperatures develop faster, mature earlier but reach smaller adult body sizes (Atkinson 1994; Ohlberger 2013). In line with TSR expectations, faster growth rates or larger size-at-age of young life stages are commonly found in both experimental, field data and modelling studies (Thresher *et al*. 2007; Neuheimer *et al*. 2011; Ohlberger *et al*. 2011; Neuheimer & Grønkjaer 2012; Baudron *et al*. 2014; Huss *et al*. 2019; van Dorst *et al*. 2019). Similarly, declines in maximum or asymptotic body size of fish have been reported to correlate with warming trends for a number of commercially exploited marine fishes (Baudron *et al*. 2014; van Rijn *et al*. 2017; Ikpewe *et al*. 2020). However, in intensively fished stocks, observed adult body sizes can decrease also for other reasons, including direct removals of large fish, or evolution towards earlier maturing and fast growth in response to fishing (Jorgensen *et al*. 2007; Audzijonyte *et al*. 2013). Moreover, decreasing adult fish size in warming waters is by far not universal. For example, no clear negative effects of warming on the body size or growth of large fish could be found in a recent experimental study (Barneche *et al*. 2019), or in a semi-controlled lake heating experiment (Huss *et al*. 2019). Similarly, across 335 coastal fish species mean species body size was similarly likely to be larger or smaller in warmer waters (Audzijonyte *et al*. 2020). Also Tu *et al*. (2018) found that temperature had a relatively minor effect on fish size structure compared to fishing, when assessing 28 stocks from the North Sea, US west coast and Eastern Bering Sea. Even when combined with fishing, only 44% of variation in size structure could be explained. Thus, the effects of temperature on body sizes may be more complex than often depicted, and we still do not fully understand the mechanisms by which temperature affects growth and body size over ontogeny (Ohlberger 2013; Audzijonyte *et al*. 2019). Increasing our understanding of these mechanisms is important because body size is a key trait in aquatic ecosystems (Andersen *et al*. 2016) and warming-induced changes in growth and size-at-age of fish populations could have implications not only for biomass and productivity, but also ecosystem structure and stability (Audzijonyte *et al*. 2013).

Physiologically structured models can address the complex interplay of direct and indirect temperature impacts on food webs, as they account for the food and size dependence of body growth through ecological interactions using bioenergetic principles. Recent applications have demonstrated decreasing maximum body sizes in fish communities due to changes in plankton abundance or size (Woodworth-Jefcoats *et al*. 2019). Similar body size responses emerge in models that focus on temperature-dependence of physiological processes, such as metabolism and feeding rates (Lefort *et al*. 2015; Guiet *et al*. 2016; Woodworth-Jefcoats *et al*. 2019), but it remains unclear to what extent these community body size shifts are driven by declining abundance of large fish versus changes in size-at-age across a range of ages.

To explore how direct and indirect effects of warming impact marine food web size structure and fisheries yields, we evaluate the impacts of temperature-driven changes in resource productivity and individual fish physiology using an example case of the Baltic Sea. The Baltic Sea constitutes a great example system, as it is a relatively well understood and species poor system (Mackenzie *et al*. 2007; Casini *et al*. 2009) that also is one of the warming hotspots globally (Belkin 2009). By using a temperature-dependent size spectrum model we analyse a set of different scenarios where either fish physiology, basal resources, or both depend on temperature, and contrast these scenarios to one another and to non-warming scenarios. We investigate the mechanisms of warming effects on body growth trajectories (size-at-age), average body sizes, population size-structure and fisheries reference points.

## Materials and Methods

In this section, we will describe the food web the model is parameterized to, the equations of the multi-species size spectrum model, how temperature dependence is implemented in the model, how the model is calibrated and lastly, how the effects of temperature are evaluated.

### Food web

We developed a multi-species size spectrum model (MSSM) (Scott *et al*. 2014), parameterized to represent a simplified version of the food web in the offshore pelagic south-central Baltic Sea ecosystem (Baltic proper) (ICES sub divisions 25-29+32, Fig. S2, *Supporting Information*) and account for temperature-dependence of processes within and between individuals (Fig. 1). This size structured food web is here characterized by three fish species: Atlantic cod (*Gadus morhua*), sprat (*Sprattus sprattus*) and herring (*Clupea harengus*), and two background resource spectra constituting food for small fish (pelagic and benthic resources). In the south-central Baltic Sea, these fish species are dominant in terms of biomass, they are the most important species commercially and they all have analytical stock assessments (ICES 2021). The pelagic background resource spectrum represents mainly phyto- and zooplankton while the benthic background resource spectrum represents benthic invertebrates, gobiidaes and small flatfish.

**Figure 1.**
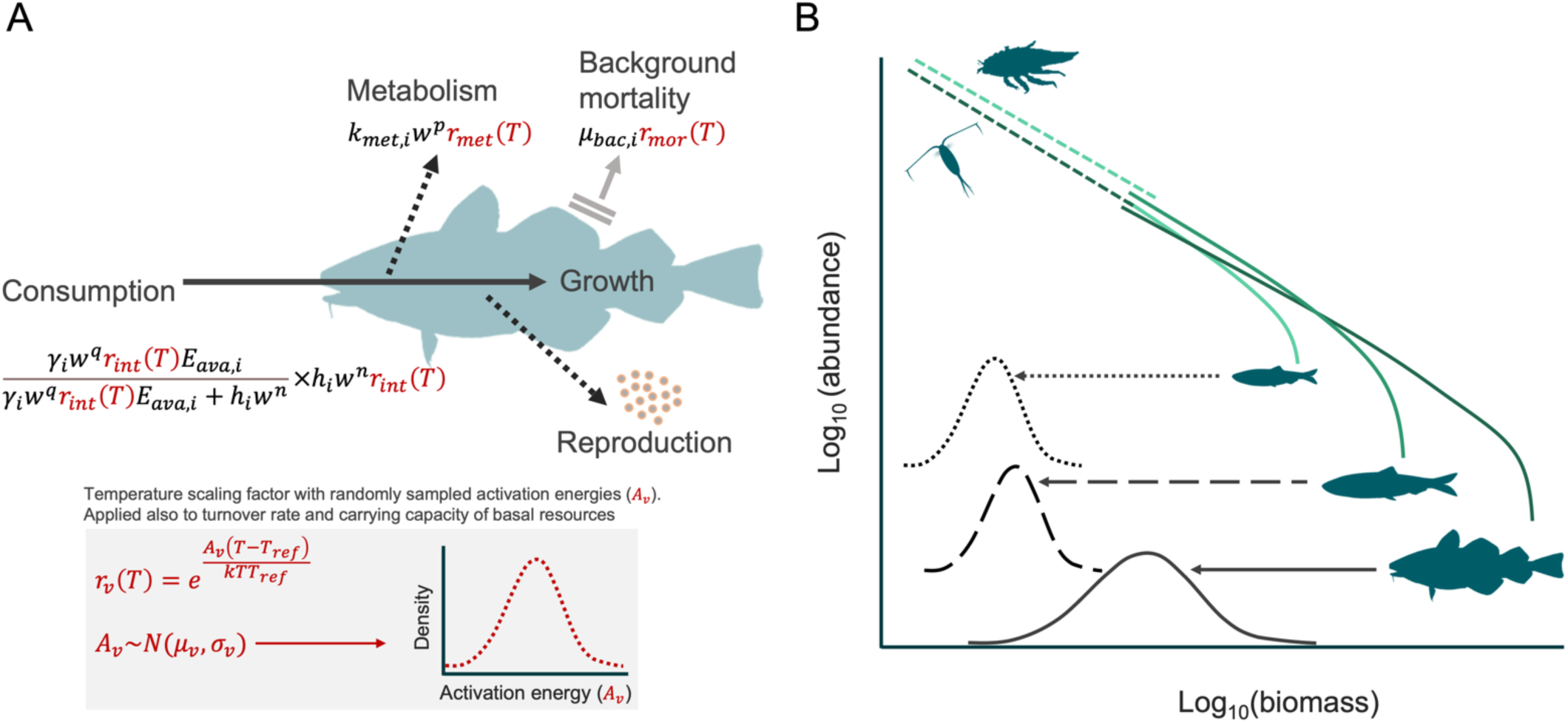
(A) Schematic representation of the individual level energy fluxes and their temperature dependence, and (B) the abundance spectrum of fishes (solid lines) emerging from food-dependent growth and mortality, and spectra of their background pelagic and benthic resource (dashed lines).

### Size spectrum model

The model is based on source code for the multi-species implementation of size spectrum models in the ‘R’-package *mizer* (v1.1) (Blanchard *et al*. 2014; Scott *et al*. 2014; R Core Team 2020), which has been extended to include multiple background resources (https://github.com/sizespectrum/mizerMR) and temperature-scaling of key physiological processes. In this section we describe the key elements of the MSSM using the same notation when possible as in previous multispecies *mizer* models for consistency (Blanchard *et al*. 2014; Scott *et al*. 2014, 2018).

In MSSMs, individuals are characterized by their weight (*w*) and species identity (*i*). The core equation is the McKendrik-von Foerster equation. Here it describes the change in abundance-at-size through time, *N*_*i*_(*w*), from food dependent somatic growth and mortality, based on bioenergetic principles (explicit modelling of energy acquisition and use):

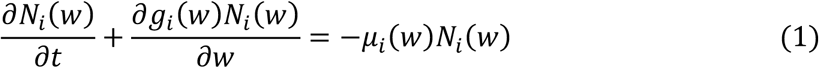

where *g*_*i*_(*w*) (g year^−1^) is somatic growth (dependent on the availability of food) and *μ*_*i*_(*w*) (year^−1^) is total mortality. At the boundary weight (*w*_0_, egg size), the influx of individuals is given by constant recruitment, which occurs at every model time step. Total mortality is the sum of the background-, starvation-, fishing-, and predation mortality. The constant species-specific allometric background mortality, *μ*_*bac,i*_, representing density- and predation independent sources of mortality, such as ageing, diseases, predation from species not included in the model, depends on the asymptotic weight of a species 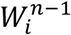 and is given by:

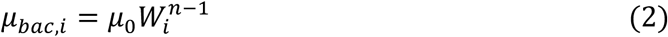

where *n* is the mass-exponent of maximum consumption rate (Hartvig *et al*. 2011) and *μ*_0_ is an allometric constant. Starvation mortality (*μ*_*stv,i*_) is assumed to be proportional to energy deficiency (defined in Eq. 11) and inversely proportional to body mass (weight, *w*), and is defined as:

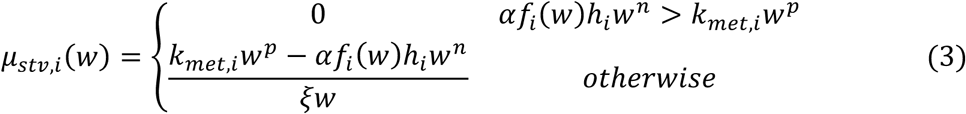

where *ξ*, the fraction of energy reserves, is 0.1 (Hartvig *et al*. 2011). Instantaneous fishing mortality (*μ*_*fis,i*_) (year^−1^) is defined as:

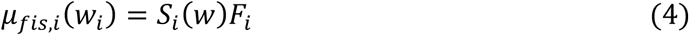

where *S*_*i*_ is the selectivity (for simplicity, we assumed knife-edge selectivity with weight at first catch corresponding to weight at maturation), and *F*_*i*_ is fishing mortality. Predation mortality (*μ*_*pre,j*_) for a prey species (or resource) *j* with weight *w*_*j*_ equals the amount consumed by predator species *i* with weight *w*_*i*_:

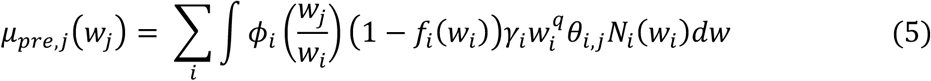

where *θ*_*i,j*_ is the non-size based preference of species *i* on species *j*, and *ϕ*_*i*_ describes the weight-based preference from the log-normal selection model (see below) (Ursin 1973). Satiation determines the feeding level *f*_*i*_(*w*) and is represented in the model with a Holling functional response type II. Satiation varies from 0 (no satiation) to 1 (full satiation):

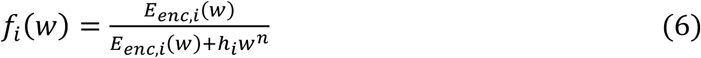

*h*_*i*_*w*^*n*^ is the allometric maximum consumption rate and *E*_*enc,i*_(*w*) is the encountered food (mass per time). The amount of encountered food for a predator of body weight *w* is given by the available food in the system multiplied with the search volume, *γ*_*i*_*w*^*q*^. Here, available food, *E*_*ava,i*_, is the integral of the biomass of all prey species (*j*) and background resources (*R*) that falls within the prey preference (*θ*_*i,j*_, *θ*_*i,R*_) and size-selectivity (*ϕ*_*i*_) of predator species *i*:

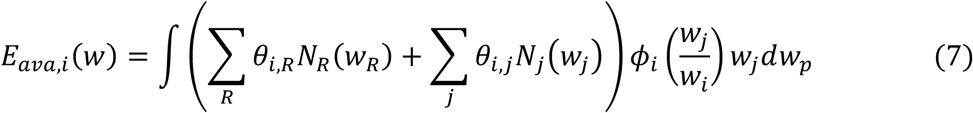

where *w*_*j*_ is the weight of prey, *θ*_*i,R*_ is the preference of species *i* for resource *R*, and *j* indicates prey (fish) species. Note that in contrast to other MSSMs (e.g., Blanchard *et al*. 2014) species in our model do not have specific preferences for other size-structured species (all values in the species interaction matrix *θ*_*i,R*_ are set to 1). However, they have different preferences for the two background resources. This helps to account for different feeding of sprat, herring and cod on benthic and pelagic resources. The size-selectivity of feeding, 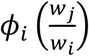, is given by a log-normal selection function (Ursin 1967):

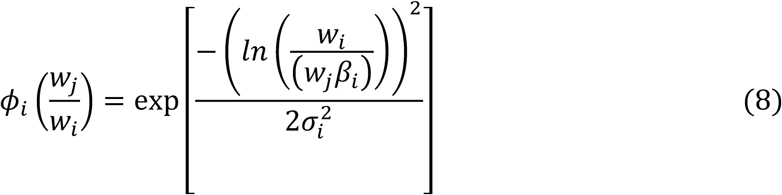

where parameters *β*_*i*_ and *σ*_*i*_ are the preferred predator-prey mass ratio and the standard deviation of the log-normal distribution, respectively. The amount of available prey of suitable sizes (Eq. 7) is multiplied with the allometric function describing the search volume (*γ*_*i*_*w*^*q*^). The allometric search volume coefficient is calculated as:

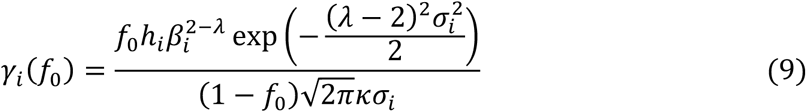

(Andersen & Beyer 2006; Scott *et al*. 2018). The actual biomass of food encountered, *E*_*enc,i*_(*w*), is defined as:

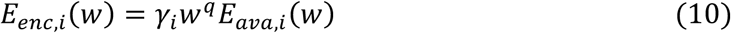

where *q* is the size-scaling exponent of the search volume. The rate at which food is consumed is given by the product *f*_*i*_(*w*)*h*_*i*_*w*^*n*^, which is assimilated with efficiency *α* and used to cover metabolic costs. Metabolic costs scale allometrically as *k*_*met,i*_*w*^*p*^. The net energy, *E*_*net,i*_, is thus:

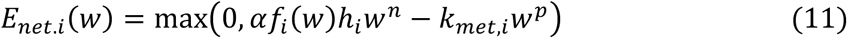

which is allocated to growth or reproduction. The allocation to reproduction (*ψ*_*i*_) increases smoothly from 0 around the weight maturation, *w*_*mat,i*_, to 1 at the asymptotic weight, *W*_*i*_, according to the function:

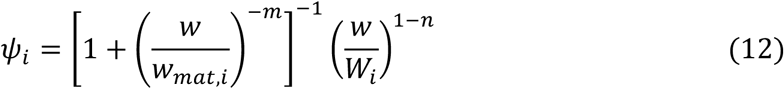

where *m* determines the steepness of the energy allocation curve, or how fast the allocation switches from growth to reproduction at around the maturation size (Andersen 2019). This function results in the growth rate, *g*_*i*_(*w*),

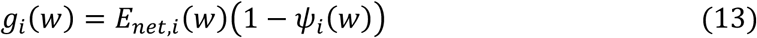

which approximates a von Bertalanffy growth curve when the feeding level is constant (Hartvig *et al*. 2011; Andersen 2019). Reproduction is given by the total egg production in numbers, which is the integral of the energy allocated to reproduction multiplied by a reproduction efficiency factor (*ϵ, erepro*) divided by the egg weight, *w*_0_, and the factor 2, assuming only females reproduce:

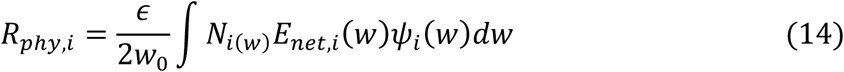

This total egg production (or physiological recruitment, *R*_*phy,i*_) results in recruits via a Beverton-Holt stock recruit relationship, such that recruitment approaches a maximum recruitment for a species *i* (*R*_*max,i*_), as the egg production increases,

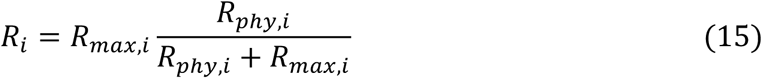

where *R*_*max,i*_ is treated as a free parameter and is estimated in the calibration process by minimizing the residual sum of squares between spawning stock biomass from stock assessments and the MSSM. The calibration also ensures that the species coexist in the model.

The temporal dynamics of the background resource (*N*_*R*_) spectra (benthic and pelagic) are defined as:

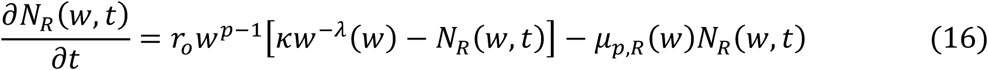

where *r*_*o*_*w*^*p*−1^ is the population regeneration rate, *κw*^−*λ*^ is the carrying capacity of the background resource and *μ*_*pre,R*_ is predation mortality on resource spectrum *R*, and *λ* is defined as −2 − *q* + *n* (Andersen 2019).

### Temperature dependence of fish physiology and resource dynamics

To study the effects of warming on the modelled ecosystem, we introduce temperature impacts on size-structured species physiological rates and background resource growth dynamics following the metabolic theory of ecology (Gillooly *et al*. 2001; Brown *et al*. 2004). Temperature affects the rate of metabolism (Clarke & Johnston 1999; Gillooly *et al*. 2001), and thus also other biological rates such as feeding and mortality (Brown *et al*. 2004; Englund *et al*. 2011; Rall *et al*. 2012; Thorson *et al*. 2017). We therefore scale rates of individual metabolism (*k*_*met*.*i*_*w*^*p*^), maximum consumption (*h*_*i*_*w*^*n*^), search volume (*γ*_*i*_*w*^*q*^) and background mortality 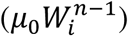 with temperature. Metabolism and consumption are key terms in the energy budget of fish (Eqns. 11-13). Thus, growth rate is not temperature-dependent directly but its relationship to temperature emerges from the temperature-scaling of metabolism and consumption. In *mizer*, metabolism represents all metabolic costs, i.e., standard, activity, and digestion. Henceforth, we assume *k*_*metei*_*w*^*p*^ scales as standard metabolic rate and refer to it as metabolism or metabolic rate. For simplicity reasons and to reduce the number of parameters and scenarios we assume that all rates scale with temperature exponentially according the Arrhenius temperature correction factor:

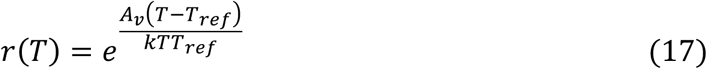

where *A*_*v*_ is the activation energy (eV) for individual rate *ν, T* is temperature (K), *T*_*ref*_ is the reference temperature (here 283.15 K, the temperature where the Arrhenius correction factor equals 1), and *k* is Boltzmann’s constant in eV K^−1^ (= 8.617 × 10^−5^ eV K^−1^). We chose an exponential temperature dependence as it provides a good statistical fit to data, is widely adopted, and because we assume that the projected change in ocean temperature in the studied time range does not lead to temperatures above physiological optima (e.g. (Righton *et al*. 2010) as an example for cod), whereafter physiological rates might be expected to decline. While temperature likely affects other physiological processes as well (such as cost of growth (Barneche *et al*. 2019) or food conversion efficiency (Handeland *et al*. 2008)), we focus on the temperature effects on metabolism, maximum consumption, search volume and mortality, as their temperature dependencies are relatively well documented (Pauly 1980; Brown *et al*. 2004; Dell *et al*. 2011; Englund *et al*. 2011; Thorson *et al*. 2017; Lindmark *et al*. 2022).

Temperature also affects plankton and benthos organisms, represented in our model through background resources. In most size spectrum models to date, climate affects primary production (and in some cases zooplankton), and this is modelled by forcing the background spectra to observed abundance-at-size of plankton from either remotely sensed variables such as chlorophyll-a, or from output (e.g., net primary production) from earth-system models (Blanchard *et al*. 2012; Barange *et al*. 2014; Jennings & Collingridge 2015; Canales *et al*. 2016; Galbraith *et al*. 2017; Reum *et al*. 2019; Woodworth-Jefcoats *et al*. 2019). These differences have been highlighted as a key source of ecosystem model uncertainties observed in global applications of size-structured models (Lotze *et al*. 2019; Heneghan *et al*. 2021). In order to isolate the effects of temperature on resource and physiological processes, we apply the temperature scaling to the terms of the background resource’s semi-chemostat growth equation (Eq. 16), i.e., their biomass regeneration rate and carrying capacity. We use the same Arrhenius correction factor with activation energy *A*_*r*_, where *r* refers to background resource parameter. We assume that as temperature goes up, the carrying capacity (*κw*^*λ*^) declines at the same rate as population regeneration (*r*_0_*w*^*p*−1^) rate increases (Savage *et al*. 2004; Gilbert *et al*. 2014), i.e. *κ* scales with temperature in proportion to 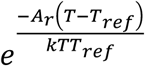. This is based on the metabolic theory of ecology (MTE), which predicts that if nutrient levels are constant, higher respiration rates lead to lower biomasses at carrying capacity (Savage *et al*. 2004; Bernhardt *et al*. 2018). To simplify the analyses, our implementation of temperature effects on the background spectrum assumes that its size structure is not affected by the temperature (the slope of the spectrum does not change)—only the overall level of background resources. As an example, using the average activation energy for resource carrying capacity (see next paragraph), the elevation of our background resource spectra (abundance at the geometric mean weight), declines with 8.7% with a 1°C increase in temperature, which is line with a previous study (Heneghan *et al*. 2019).

Activation energies are estimated with uncertainty and they vary substantially between processes, species, and taxonomic groups. To account for this uncertainty, here we parameterized 200 projections of the food web model using randomly sampled activation energies from normal distributions with rate-specific means and standard deviations. For metabolism and maximum consumption, we acquired means and standard deviations of posterior distributions provided in (Lindmark *et al*. 2022). For search volume we assumed that it scales identically to maximum consumption, because both rates are related to feeding processes. For background mortality we assumed identical scaling to metabolism, as longevity is linked to life span and metabolic rate (Brown *et al*. 2004; McCoy & Gillooly 2008; Munch & Salinas 2009). For background resource activation energies, we use the point estimate of the activation energy (slope from a linear regression of natural log of growth rate as a function of Arrhenius temperature (1/*kT* [eV^−1^])) from experimental data in Savage *et al*. (2004) as the mean. These data consisted of protists, algae and zooplankton, and were extracted using the software WebPlotDigitizer v. 4.1 (Rohatgi 2012). The standard deviation was approximated by finding the value that resulted in 95% of the normal distribution being within the confidence interval of the linear regression. For each of the 200 parameter combinations, each of the six rate activation energy parameters was sampled independently from their respective distribution and the model was projected to 2050. We then quantified the overall mean response and the ranges of predictions resulting from 200 randomly parameterised simulations and visualized it for the analysis of growth (size-at-age) and mean size.

We acknowledge that these scenarios are very simplified for evaluating changes in resource productivity versus physiology with warming, and that they do not necessarily reflect the predicted conditions in the Baltic Sea, nor all the potential pathways by which climate changes affects the environmental conditions in the Baltic Sea. However, the simplicity allows us to contrast effects of warming on basal food resources versus individual physiology of fish.

### Model calibration

Here we present a summary of the calibration approach—a more detailed description of the step-by-step calibration protocol can be found in *Model calibration and validation, Supporting Information*. The model was calibrated to average spawning stock biomasses (*SSB*_*i*_) from stock assessment data for cod, herring and sprat (ICES 2013, 2015) in 1992-2002, using average fishing mortalities (*F*_*i*_) in the same time frame. Ideally, the period for calibration should exhibit relative stability, but such periods do not exist in the Baltic Sea, which is greatly influenced by anthropogenic activities and has undergone dramatic structural changes over the last four decades (Möllmann *et al*. 2009). We chose to calibrate our model to the time period of 1992-2002 as in Jacobsen *et al*. (2017), which is a period after an ecological regime shift, characterized by high fishing mortality on cod, low cod and herring abundance and high sprat abundance (Gårdmark *et al*. 2015) (Fig. S4). The cut-off at 2002 also ensured that we did not calibrate the model to the period starting from mid 2000’s when the growth capacity, condition, proportion of large fish in the population, and reproductive capacity of cod started to decline rapidly (Svedäng & Hornborg 2014; Casini *et al*. 2016; Mion *et al*. 2018, 2021; Neuenfeldt *et al*. 2020).

Model calibration was done by tuning the maximum recruitment parameter *R*_*max,i*_ for the three fish species while holding temperatures at *T*_*ref*_. *R*_*max,i*_ determines the maximum number of offspring that can be produced by a population in a given time step and serves as a density independent cap on reproduction. This parameter determines how species will respond to exploitation and perturbations, and is one of the main parameters that is calibrated in multi-species models (e.g., Blanchard *et al*. 2014; Jacobsen *et al*. 2017). We used the “L-BFGS-B” algorithm (Byrd *et al*. 1995) in the ‘R’-optimization function ‘*optim*’ to minimize the residual sum of squares between the natural log of spawning stock biomass estimated in stock assessment output (ICES 2013, 2015) and those emergent in the model for the years 1992-2002. The optimization procedure resulted in close agreement between SSB from the model and from stock assessments in the calibration time frame. Projections from 1992 and 2012 also generally tracked the assessment SSBs (correlation coefficients of 0.65, 0.94 and 0.54 for cod, herring and sprat, respectively). However, hindcasts (1974-2012) revealed no correlation between assessment SSB and the model, while for herring and cod the general trends were captured relatively well. The low ability to reconstruct historical biomasses is likely due to the regime shift occurring between 1988-1993 (Möllmann et al. 2009). Growth curves emerging from the model were in close agreement with von Bertalanffy curves fitted to length-at-age data from trawl surveys (Fig. S6), after a stepwise manual increase of the constant in the allometric maximum-consumption rate (*h*_*i*_) (*Supporting Information*). The level of density dependence imposed by the stock-recruitment function (see Eq. 14-15) was also evaluated by assessing the ratio of the physiological recruitment, *R*_*phy,i*_, to the recruitment *R*_*i*_ (Jacobsen *et al*. 2017) (*Supporting Information*). These final values mean that stock recruitment is sensitive to the stock biomass, but there is some density dependence limiting recruitment (i.e., not all spawn produced become recruits). The fishing mortality leading to the highest long-term yield (*F*_*MSY*_) from the model (estimated for one species at the time while keeping each species at their mean assessment *F*_*MSY*_) were in agreement with the assessment *F*_*MSY*_ for sprat and herring. For cod, *F*_*MSY*_ is lower in the size-spectrum model than in stock assessments.

### Analysis of responses to warming

We analyse the effects of warming on the size-structure food web in two different ways: by projecting the food web to 2050 with time-varying sea surface trends, and by projecting the model for 200 years with fixed temperatures above or below *T*_*ref*_. The first set of simulations aimed to assess possible fish population responses to the expected temperature changes, while the second was aimed at exploring effects of temperature on fisheries yield and *F*_*MSY*_ at steady state.

For the time-varying simulations, models were projected with historical annual fishing mortalities (1974-2012) (ICES 2013, 2015) and sea surface temperature trends (1970-2050, acquired from the regional coupled model system RCA4-NEMO under the RCP 8.5 scenario) (Dieterich *et al*. 2019; Gröger *et al*. 2019). These relative temperature trends (relative to mean in 1970-1999) are scaled by adding a constant so that the average temperature in the in the calibration time period is *T*_*ref*_ (10°C). To ensure steady state was reached before time-varying fishing mortality and temperature was introduced (1974 and 1970, respectively), we applied a 100-year burn-in period using the first fishing mortality and temperature value in the respective time series (Fig. S12). For each species, we used the fishing mortality at maximum long-term (‘sustainable’) yield, *F*_*MSY*_ in the years 2012-2050 (Fig. S12). These were derived from the size spectrum model by finding the fishing mortalities resulting in highest yields at *T*_*ref*_ (Fig. S9). We evaluated the effects of warming on weight-at-age, population mean weight and abundance-at-weight for each species. This was done for both absolute values, and by comparing warming food webs in 2050 to a baseline scenario where no warming occurred post 1997 (the mid-point of calibration time window, where temperature averages *T*_*ref*_) (Fig. S12). In this way the three scenarios considered contrast the effects of temperature affecting fish physiology, their resources or both.

For the non-time varying temperature projections we specified a range of constant (not time-varying) temperatures and fishing mortalities, expressed as proportions of *T*_*ref*_ and *F*_*MSY*_ at the reference temperature 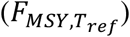, respectively, and projected the models to steady state (200 years). We explored scenarios were temperature ranged between 0.75 to 1.25 of *T*_*ref*_, and *F*_*MSY*_ ranged between 0.1 and 2 of 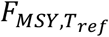 With the full factorial combination of these scenarios, this gave a total of 1989 scenarios. These simulations were done to explore the effect of temperature on fisheries yield and *F*_*MSY*_.

## Results

### Effects of warming on size-at-age depend on physiological temperature-dependence

The inclusion of temperature effects on fish physiological processes has a strong influence on the projected size-at-age in 2050 under the RCP 8.5 emission scenario, relative to the baseline projection (no warming) (Fig. 2). Temperature-dependence of feeding rates have a particularly large effect (Fig. S15). Warming positively affects size-at-age when temperature affected metabolism, maximum consumption, search volume and mortality, regardless of whether temperature impacted background resource dynamics (Fig. 2). In contrast, the scenarios without temperature-dependent physiological processes all lead to size-at-age decreasing with warming (Fig. 2). In scenarios with temperature-dependent physiological processes, the effects on size-at-age are positive and declines with age. When only resources are affected by temperature, small individuals have the largest relative decrease in size-at-age, and this negative effect of warming declines with age (Fig. 2).

**Figure 2.**
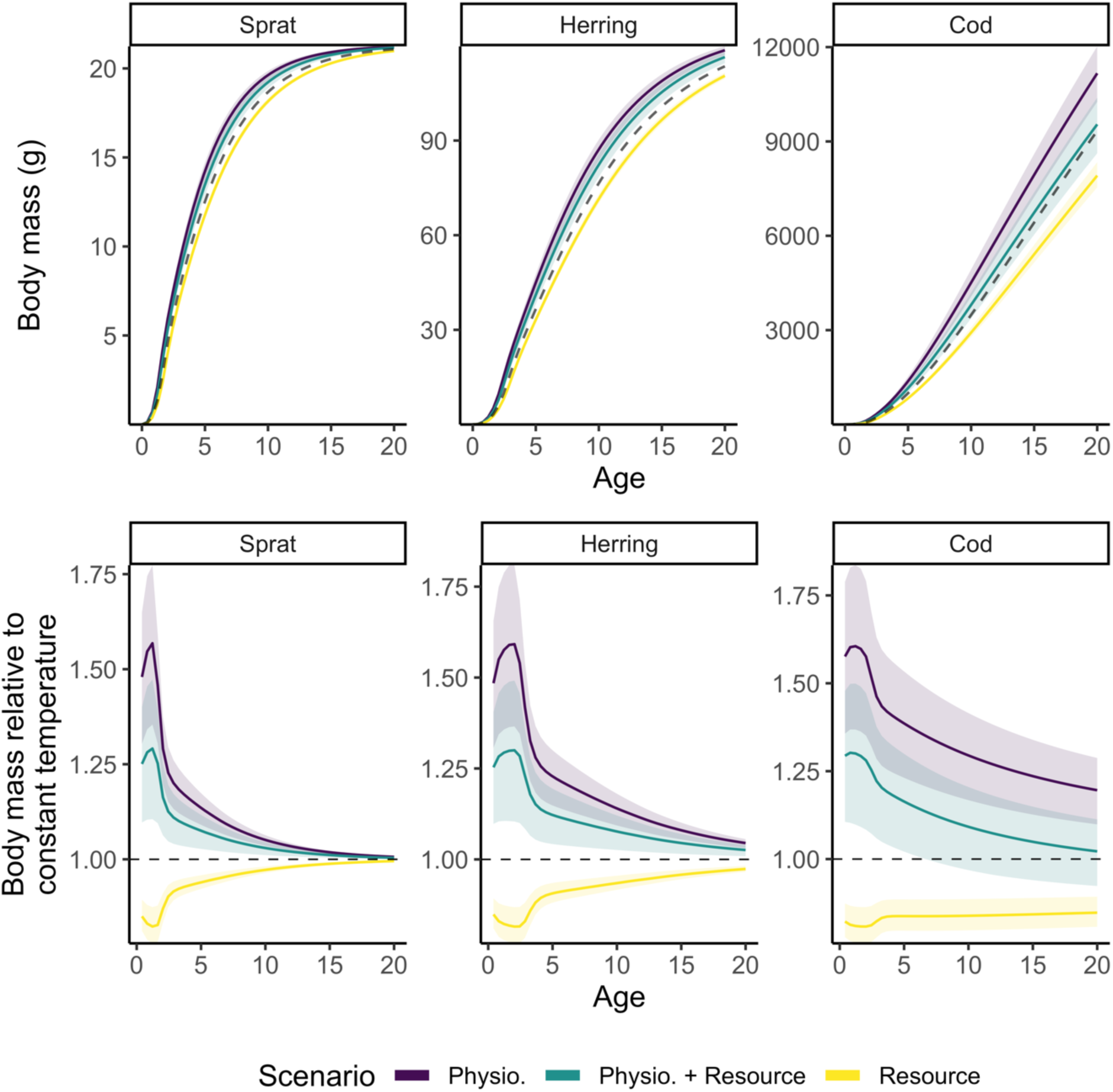
Individual growth trajectories of sprat, herring, and cod from model projections to year 2050 assuming warming according to RCP 8.5 while keeping fishing mortality at F_MSY_ levels from the size spectrum model. Top row shows size-at-age and bottom row shows size-at-age relative to a non-warming scenario. The dashed line in the top row depicts projections assuming a non-warming scenario and thus constitutes a baseline prediction. Colours indicate different temperature-scaling scenarios. Shaded areas encompass the 2.5 and 97.5 percentiles from the set of 200 simulations with randomly assigned activation energies.

Despite the relatively narrow range of activation energies for physiological rates considered here (Fig. S3; Table S3), the uncertainty in projected size-at-age associated with variation in the activation energies is large (Fig. 2). In the scenario where both physiology and resources are affected by temperature, the range of predicted changes in size-at-age vary at approximately +10% to +40% (Fig. 2). These changes in size-at-age seem to be driven by the temperature-dependence of maximum consumption rate (*h*_*i*_*w*^*n*^(*T*)) increasing the actual consumption rates (*f*_*i*_(*w*)*h*_*i*_*w*^*n*^(*T*)), but having almost no effect on the feeding or satiation levels (Eq. 6; Fig. S13).

### Fewer large individuals cause reductions in mean population body size

Increases in size-at-age (Fig. 2) do not always lead to increased mean body size in the populations (Fig. 3), due to changes in the relative abundances at size and in this way shifting population size structure (Fig. 4). Changes in the size-structure varied across species without a clear and consistent pattern across species and scenarios.

**Figure 3.**
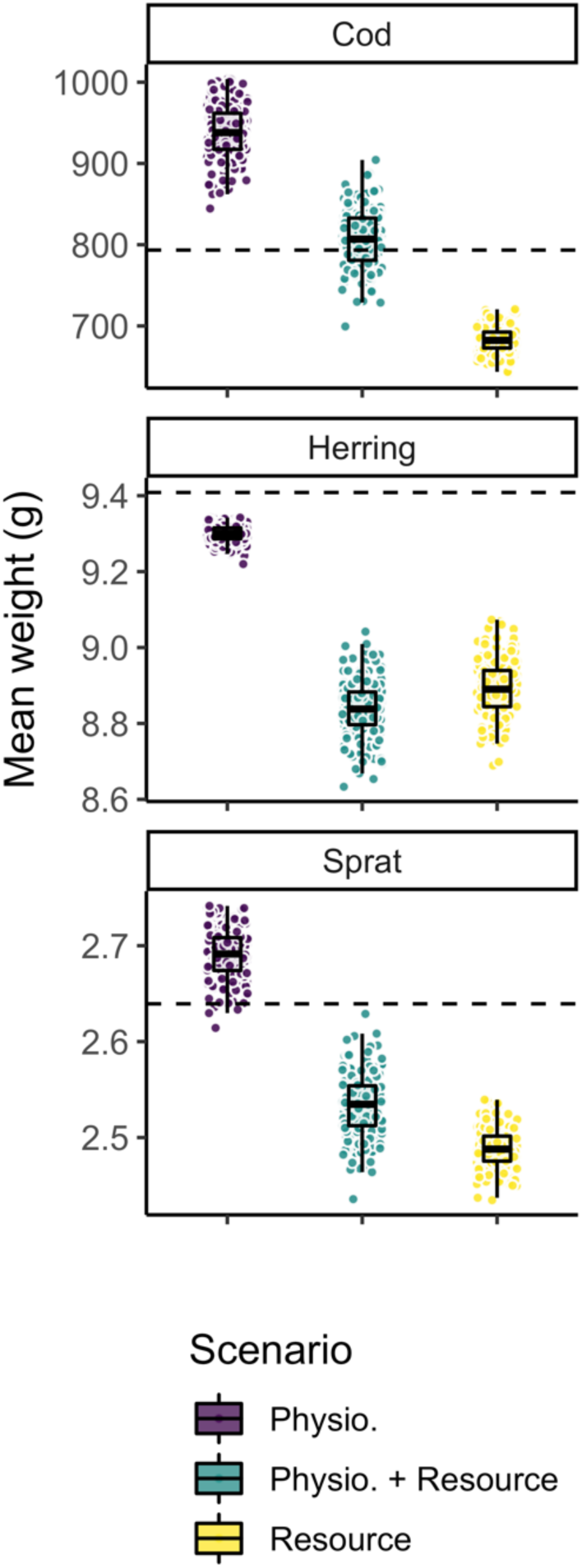
Mean weight across all individuals in the populations of sprat, herring and cod from model projections to year 2050 assuming warming according to RCP 8.5 while keeping fishing mortality at F_MSY_ levels from the size spectrum model. The dashed horizontal line depicts projections assuming no temperature increase and thus constitutes a baseline prediction. Each dot represents one of the 200 simulations, each with randomly assigned activation energies. Boxplots depict 25%, 50% and 75% quantiles of the 200 simulations in each scenario.

**Figure 4.**
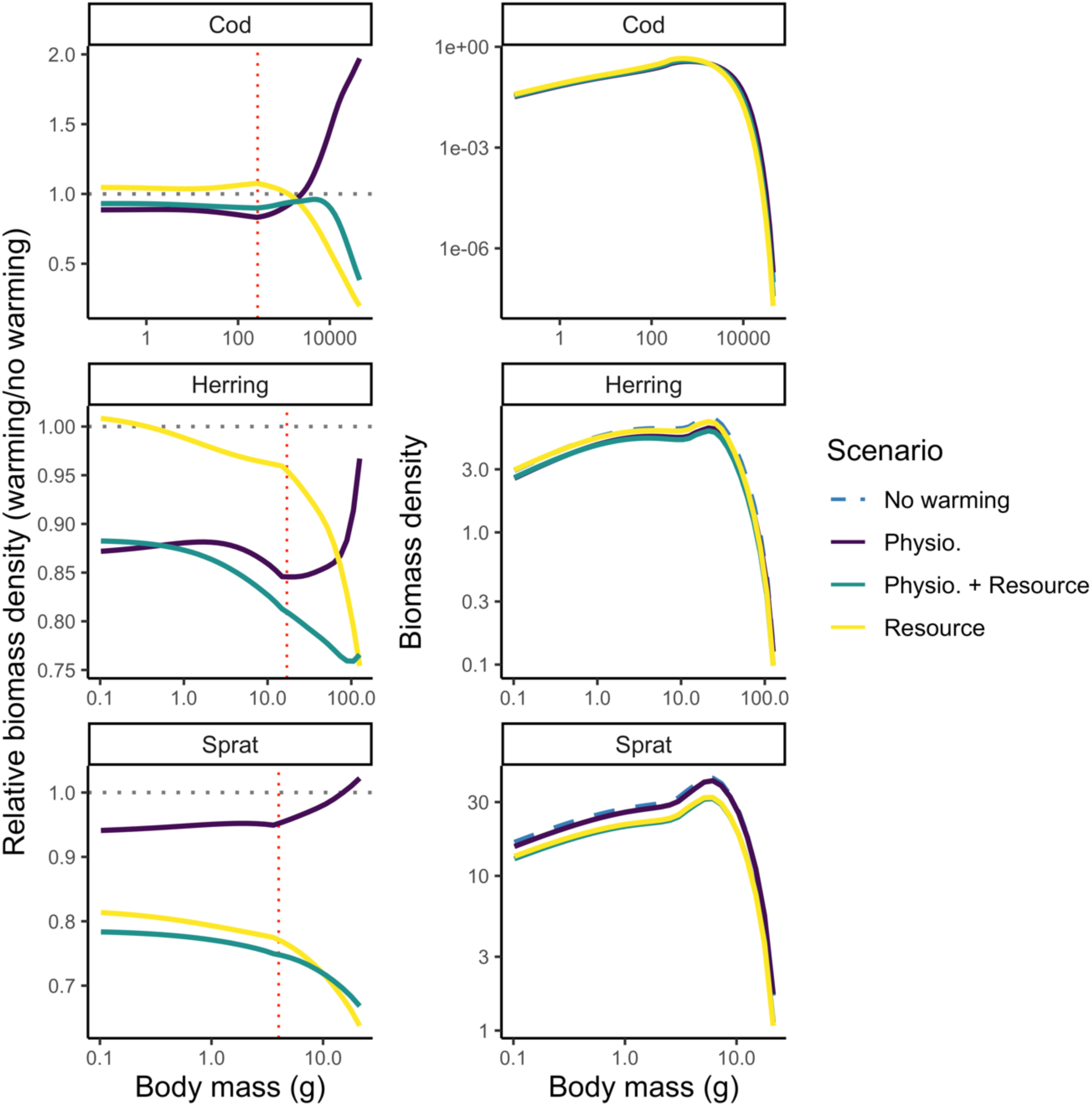
Projected abundance-at-weight by species for different scenarios of temperature scaling indicated by colours (and line types in the right column due to overplotting) in 2050 assuming fishing mortality held at F_MSY_ levels from the size spectrum model. The left column shows abundance-at-weight relative to a non-warming scenario and the right column shows absolute abundance-at-weight with the non-warming scenario shown in black. Vertical red dotted line indicates weight-at-maturation and horizontal black dotted lines indicate the baseline projection (no warming). Only mean activation energies are used (Table S3, Supporting Information).

The only scenario where mean body weight on average increases is where temperature only affects physiology and not the resource (Fig. 3). In such cases, body weight increases with warming, but only for cod and sprat. For cod this increase is strong and is driven by both faster growth rates (larger size-at-age) and large increases in the abundance of large fish (∼10 kg) (Figs. 2, 4). For sprat the mean body weight in the populations increased only marginally and is mostly driven by faster growth rates and increased relative abundance of fish above 10 g (Figs. 2, 4). In contrast, scenarios where only resources are affected by temperature, relative numbers of large individuals and therefore mean body size of all species goes down. For herring, all scenarios lead to smaller mean body sizes in the population, and the relative (to non-warming simulation) abundance-at-weight declines with mass in most of the size range, with increases only in the very smallest size classes (< 1g; Fig. 4).

### Temperature and fishing: higher sustained exploitation rates but reduced yields in warmer environments

Our simulations applying a range of stable (not time-varying) temperature and fishing scenarios showed that warming led to higher or equal *F*_*MSY*_ (i.e., the fishing mortality leading to maximum sustainable yield), but lower yields (Fig. 5-6). *F*_*MSY*_ declines with warming for herring when only resources are temperature dependent, and *F*_*MSY*_ for sprat declines resources are temperature dependent, else *F*_*MSY*_ increases. Yields however, decline for all species in all scenarios except for cod when only physiological processes are temperature dependent. The increase in *F*_*MSY*_ is likely due to the enhanced growth rates, which allow higher fishing mortalities without impairing population growth. Cod in the scenario with only physiological scaling is the exemption. The model projects higher yields as temperature increase, due to the increase in growth rate, average size and relative abundance of large individuals (See Figs. 2-4). In general, the highest relative yield is found at the coolest temperatures and *F* slightly lower than *F*_*MSY*_ at the reference temperature (Fig. 6). The decline in relative yields of herring and sprat in all scenarios (Fig. 5) is likely driven by the warming-induced decline in abundance, due to resource limitation (Fig. 4).

**Figure 5.**
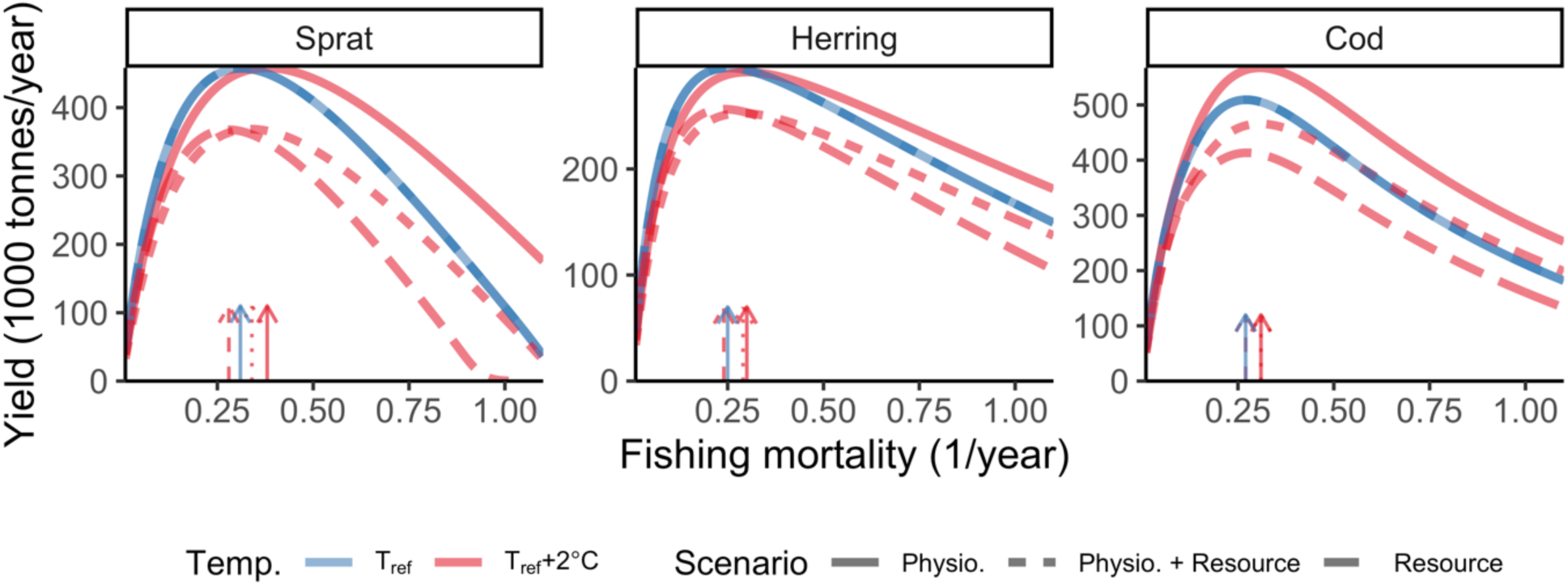
Steady state biomass yield assuming knife edge selectivity at maturation size under two constant temperature simulations and three scenarios for temperature dependence. Colours indicate temperature, where blue means T = T_ref_ (i.e., no temperature effects), and red depicts warm temperature, here T = T_ref_ +2°C. Dashed lines correspond to resource dynamics being temperature dependent, dotted lines correspond to physiological rates and resource dynamics being temperature dependent and solid lines depicts only physiological temperature scaling. Arrows indicate fishing mortality (F) that leads to maximum sustainable yield (F_MSY_). F is held constant at the mean F during calibration (mean 1992-2002) for the two other species while estimating yield curves for one species. Note the different scales between species. Only mean activation energies are used (Table S3, Supporting Information).

**Figure 6.**
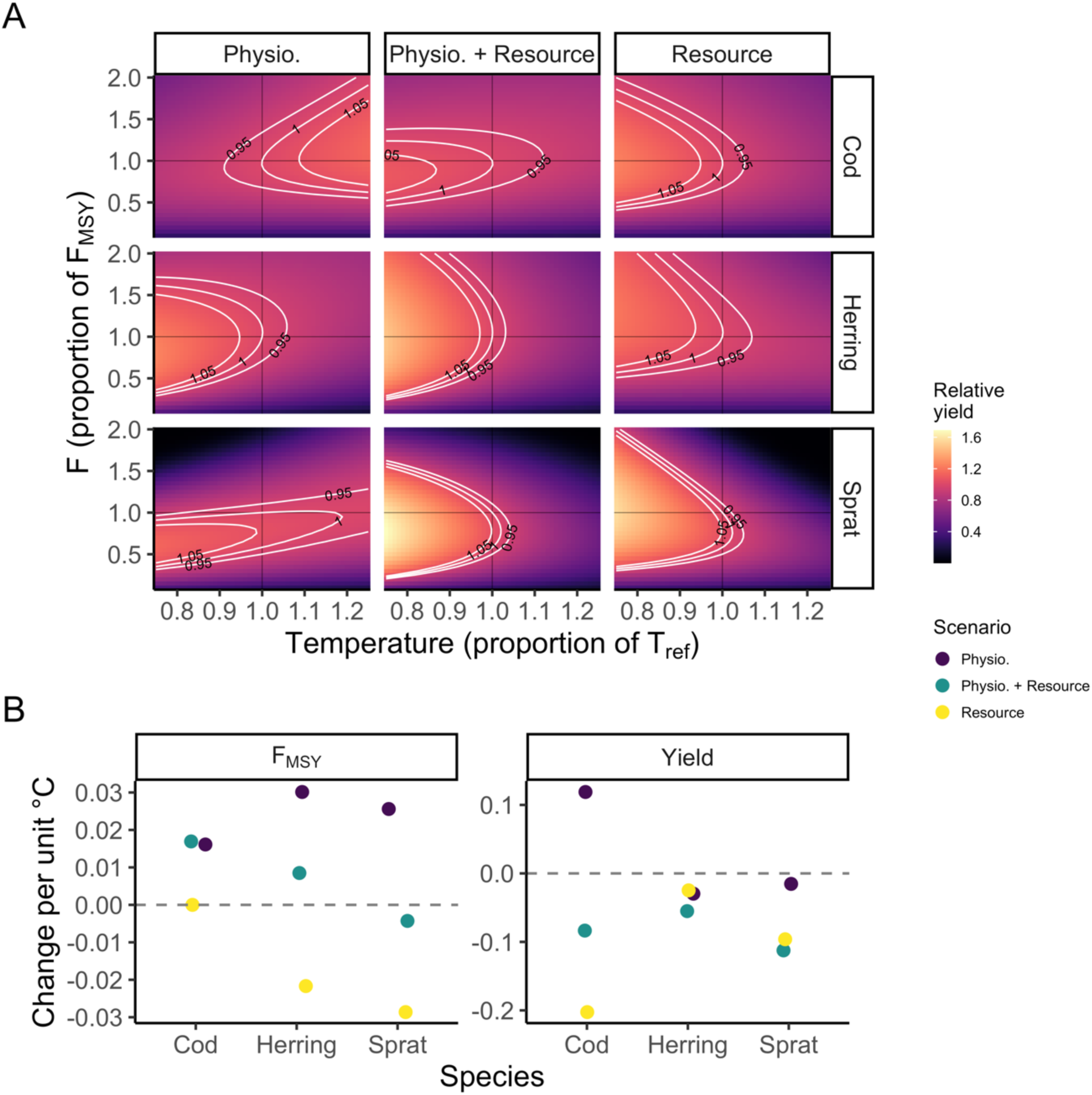
Example of fisheries yield (A) at steady state relative to MSY at T_ref_ (no effect of temperature) from simulations with constant (not time-varying) temperatures with the three temperature dependence scenarios (columns), and how F_MSY_ and yields change with temperature. In panel A, The y-axis shows fishing mortality, F, as a proportion to F_MSY_ (as estimated from the size spectrum model) at T_ref_ and the x-axis shows temperature as a proportion of T_ref_. The other two species are held at their F_MSY_ when one species’ F is varied. Panel B shows slopes of F_MSY_ (left) and yield (right) over temperature. Only mean activation energies are used (Table S3, Supporting Information).

## Discussion

### Combined temperature impacts on fish growth rates, body size and fisheries yield

Using a size-structured and species-resolved food web model, we demonstrate how climate warming affects growth rates (size-at-age), population mean size and size-structure of interacting exploited fish species and assess its implications for fisheries yield. To do so, we contrasted the effects of warming on resource productivity and individual level physiology (metabolism, feeding and background mortality) of fish. We found that warming leads to increased size-at-age of fishes when temperature-dependence is included in physiological rates. This effect is strongest in juveniles of all three fish species. Though, despite increased growth rates, in most cases, warming leads to smaller mean body size in the population, lower spawning stock biomass (biomass of mature fish) and reduced yields. When temperature affects only the background resource species, the size-at-age declines for fish of all sizes.

Mechanistic that models explore warming-driven declines in community-wide average body size often find these declines to be driven by lower food abundance or decreased energy transfer efficiency in the food web, due to a combination of declines in plankton density and shifts towards dominance of smaller plankton at higher temperatures (Lefort *et al*. 2015; Woodworth-Jefcoats *et al*. 2015, 2019). This leads to a community wide decline in mean size of fish, where large bodied species become relatively fewer. The cause of these community-level changes are different from those expected at an individual species level, where temperature can either lead to size-at-age changes over ontogeny (in accordance with the temperature-size rule), or a change in the relative abundance of small vs large individuals. TSR (the temperature-size rule) predicts higher growth rates and thus size-at-age of juveniles, but smaller adults body sizes (Atkinson 1994), although the physiological processes that lead to these changes remain debated (Audzijonyte *et al*. 2019). In our model, we include scenarios that reflect both warmer temperatures impact on food abundance as well physiological changes in metabolism and food intake rates. Scenarios with only temperature dependence of resource dynamics lead to declines in size-at-age (that in addition were strongest in young fish). This does not match general observations and predictions of how body growth is affected by warming (Thresher *et al*. 2007; Morita *et al*. 2010; Huss *et al*. 2019; Lindmark *et al*. 2022), and is not in accordance with the TSR. In contrast, inclusion of physiological temperature dependence leads to projections more in line with general observations from field data, which often find increased size-at-age that is strongest and positive for small individuals, and that this effect diminishes over ontogeny (Thresher *et al*. 2007; Huss *et al*. 2019).

The increase in body growth that we in find is in general not sufficient for maintaining similar mean population body sizes and size-structure, if resource carrying capacities decline with warming, because this reduces the relative abundance of large fish. Mean body size in the population and yields therefore decline in the scenario with temperature dependence of both resource dynamics and physiology. These predictions on the net effect of warming are in line with similar models using empirically derived static plankton spectra (Blanchard *et al*. 2012; Canales *et al*. 2016; Woodworth-Jefcoats *et al*. 2019), and empirical studies (van Dorst *et al*. 2019). If, however, resource carrying capacity would not decline with temperature, our results show that the increased body growth potential in fish due to higher metabolic and feeding rates can lead to changes towards dominance of larger fish in some populations. This is important to consider, given that predictions about effects of climate change on primary production are uncertain and show large regional variability (Steinacher *et al*. 2010). These results show that it is important to account for both direct (physiology) and indirect (resources) effects of temperature in order to explain results such as increased growth rates and size-at-age but overall smaller-bodied populations, as also found in other studies (Ohlberger *et al*. 2011; Ohlberger 2013; Neubauer & Andersen 2019; Gårdmark & Huss 2020).

In fisheries stock assessment, plastic body growth was generally thought to be less important for stock dynamics than environmentally driven recruitment variation, density dependence at early life stages and mortality (Hilborn & Walters 1992; Lorenzen 2016). Due to the accumulating evidence of time-varying and climate-driven changes in vital rates (survival, growth and reproduction), their relative importance for fisheries reference points and targets are now becoming acknowledged (Thorson *et al*. 2015; Lorenzen 2016). In our modelling system, we find that maximum sustainable yields (*MSY*) and the fishing mortality leading to *MSY*, i.e., *F*_*MSY*_, vary with both temperature and between modelling scenarios, and largely depends on the net effect of temperature on abundance-at-size and body growth rates. When temperature affects both the background resources (mainly declining carrying capacity) and fish physiology, warming tends to increase *F*_*MSY*_, but the yield (*MSY*) derived at this exploitation rate is lower. The decline in yields with warming is due to reduced resource availability, lowering overall fish abundance, and is in line with earlier studies (Blanchard *et al*. 2012; Lotze *et al*. 2019). In addition, the warming-induced decline in relative abundance of fish above minimum size caught in fisheries further decreases yields in our model. At the same time, higher growth rates (size-at-age), occurring when temperature affects metabolism and intake rates in particular, can cause *F*_*MSY*_ to increase with warming (Thorson *et al*. 2015). These reference levels should not be viewed as absolute reference points, and the specific results may depend on the model calibration procedure. However, our findings suggest that climate change predictions on fisheries productivity must consider both temperature impacts on vital rates, in particular body growth, as well as bottom-up processes and their effects on both the overall abundance and size-structure of the stock. It also indicates that because productivity may decline with warming in large parts of the oceans (Kwiatkowski *et al*. 2020) (although there is large variation in these predictions across ecosystems (Steinacher *et al*. 2010)), reduced fisheries yields may be common in a warming world.

### Parameterizing and modelling temperature effects

Including physiological temperature-dependence can strongly influence predictions of warming-effects and it allows for detailed understanding of temperature effects on populations and food webs via both individual bioenergetics and the emerging responses in fish body growth rates. However, it also requires more parameters, which in turn may vary across species. This could reduce generality of predictions and increased challenges in parameterizing models of data poor systems. We approached this by applying random parameterization, rather than fixed values of temperature dependence. To capture the uncertainty of our approach, we sampled parameters from distributions based on estimates of activation energies of physiological rates in the literature (Lindmark *et al*. 2022). This approach revealed that in terms of body growth and mean body size in populations, the combination of activation energies can determine whether the mean size increases or decreases with warming, and at what age body sizes decline relative to the current temperatures (degree of decline in size-at-age). Hence, better knowledge of the temperature-dependence of rates of biological processes is needed and these parameters should be chosen carefully, and their uncertainty acknowledged in future modelling studies.

To disentangle temperature effects on background resources and physiological processes, we modelled temperature dependence of resources by scaling their parameters with the same general Arrhenius equation (Gillooly *et al*. 2001) that we used to scale the physiological processes in fish. Other similar studies that use size spectrum models with physiological temperature-dependence instead import the plankton spectra from climate and earth systems models (Woodworth-Jefcoats *et al*. 2019) or from satellite data (Canales *et al*. 2016). Such approaches may lead to predictions that are more relevant for a specific system. However, it also becomes more difficult to separate the mechanisms behind the observed changes, as the resource dynamics then are externally forced and cannot respond to changes in the modelled food web. Moreover, populating a resource size spectrum based on observed data can be difficult as observed spectra result from both predation and bottom-up processes. As an alternative, our approach of directly scaling the carrying capacity or turnover rates of background resources with temperature provides a coherent way to model temperature-dependencies across trophic levels. The resource dynamics are then impacted by any warming-driven changes in predators, as well as inherent temperature-dependent dynamics, rather than driven by external data (Canales *et al*. 2016) or models (e.g., Woodworth-Jefcoats *et al*. 2019). On the downside, this approach means relying on many major simplifications with respect to resource dynamics. In addition, our scenarios only include identical temperature dependencies and baseline carrying capacity of pelagic and benthic resources, and only negative effects of temperature on resource carrying capacity. These may reflect the global decline in primary production (Steinacher *et al*. 2010) commonly predicted by coupled climate models. It would be straightforward to model increases in carrying capacity with our approach by using positive activation energies. It is also possible to include temperature-effects of the slope of the size spectrum, as this is often found to be negatively related to temperature (e.g., (Morán *et al*. 2010; Yvon-Durocher *et al*. 2011; Canales *et al*. 2016; Woodworth-Jefcoats *et al*. 2019), but see also Barnes *et al*. (2011) for a non-significant negative effect on the size-spectrum slope).

## Conclusion

Ecological forecasting is inherently difficult, and climate change alters the already complex causal pathways that drive ecosystem dynamics. Size spectrum models have successfully been used to evaluate size-based mechanisms and structuring forces in ecosystems (Andersen & Pedersen 2009; Szuwalski *et al*. 2017; Reum *et al*. 2019). In this study, we have highlighted the important role of temperature-dependent individual-level metabolism and feeding rates for emerging size-at-age patterns that are in line with general observations and predictions (e.g., with the TSR). These also affect the levels of exploitation that leads to maximum sustainable yields, and the corresponding yields. Hence, accounting for temperature-dependence of both ecological and physiological processes underlying population dynamics is important for increasing our understanding of how and by which processes climate change affects individuals in food webs and resulting effects on fisheries yields, which is needed to generalize across systems and into novel conditions.

## Acknowledgements

Thanks to Romain Forestier and Jonatan Reum for contributing to developing code on temperature-dependence in *mizer* during a workshop, Ken Haste Andersen for helpful discussion on model calibration, Christian Dietrich for providing temperature data. We also thank two anonymous reviewers for feedback that improved the manuscript, ICES staff and all involved in all stages of data collections, the helpful *mizer* community, Elizabeth Duskey and Magnus Huss for providing useful input. This study was supported by grants from the Swedish Research Council FORMAS (no. 217-2013-1315) and the Swedish Research Council (no. 2015-03752) (both to AG).

## Author contributions

The code was first developed from *mizer* (Scott *et al*. 2019) by AA to include multiple background resources, all authors contributed to developing the code to include temperature. ML conceived the idea. All authors contributed to study design. ML parameterized the model with input from AG. ML performed analysis and wrote the first draft. All authors contributed to writing the paper and to revisions.

## Data availability

All model code (parameterization, calibration and analysis) and data are available on GitHub (https://github.com/maxlindmark/mizer-rewiring/tree/rewire-temp/baltic) and will be deposited on Zenodo upon publication.

## References

Andersen, K.H. (2019). Fish Ecology, Evolution, and Exploitation: A New Theoretical Synthesis. Princeton University Press.

Andersen, K.H., Berge, T., Gonçalves, R.J., Hartvig, M., Heuschele, J., Hylander, S., et al. (2016). Characteristic Sizes of Life in the Oceans, from Bacteria to Whales. Ann Rev Mar Sci, 8, 217–241.

Andersen, K.H. & Beyer, J.E. (2006). Asymptotic Size Determines Species Abundance in the Marine Size Spectrum. The American Naturalist, 168, 8.

Andersen, K.Haste. & Pedersen, M. (2009). Damped trophic cascades driven by fishing in model marine ecosystems. Proceedings of the Royal Society of London B: Biological Sciences, 277, 795–802.

Atkinson, D. (1994). Temperature and organism size—A biological law for ectotherms? In: Advances in Ecological Research. Elsevier, pp. 1–58.

Audzijonyte, A., Barneche, D.R., Baudron, A.R., Belmaker, J., Clark, T.D., Marshall, C.T., et al. (2019). Is oxygen limitation in warming waters a valid mechanism to explain decreased body sizes in aquatic ectotherms? Global Ecology and Biogeography, 28, 64–77.

Audzijonyte, A., Kuparinen, A., Gorton, R. & Fulton, E.A. (2013). Ecological consequences of body size decline in harvested fish species: positive feedback loops in trophic interactions amplify human impact. Biology Letters, 9, 20121103.

Audzijonyte, A., Richards, S.A., Stuart-Smith, R.D., Pecl, G., Edgar, G.J., Barrett, N.S., et al. (2020). Fish body sizes change with temperature but not all species shrink with warming. Nat Ecol Evol, 4, 809–814.

Barange, M., Merino, G., Blanchard, J.L., Scholtens, J., Harle, J., Allison, E.H., et al. (2014). Impacts of climate change on marine ecosystem production in societies dependent on fisheries. Nature Clim Change, 4, 211–216.

Barneche, D.R., Jahn, M. & Seebacher, F. (2019). Warming increases the cost of growth in a model vertebrate. Functional Ecology, 33, 1256–1266.

Barnes, C., Irigoien, X., De Oliveira, J.A.A., Maxwell, D. & Jennings, S. (2011). Predicting marine phytoplankton community size structure from empirical relationships with remotely sensed variables. J Plankton Res, 33, 13–24.

Baudron, A.R., Needle, C.L., Rijnsdorp, A.D. & Marshall, C.T. (2014). Warming temperatures and smaller body sizes: synchronous changes in growth of North Sea fishes. Global Change Biology, 20, 1023–1031.

Belkin, I.M. (2009). Rapid warming of large marine ecosystems. Progress in Oceanography, 81, 207–213.

Bernhardt, J.R., Sunday, J.M. & O’Connor, M.I. (2018). Metabolic Theory and the Temperature-Size Rule Explain the Temperature Dependence of Population Carrying Capacity. The American Naturalist, 192, 687–697.

Blanchard, J.L., Andersen, K.H., Scott, F., Hintzen, N.T., Piet, G. & Jennings, S. (2014). Evaluating targets and trade-offs among fisheries and conservation objectives using a multispecies size spectrum model. Journal of Applied Ecology, 51, 612–622.

Blanchard, J.L., Jennings, S., Holmes, R., Harle, J., Merino, G., Allen, J.I., et al. (2012). Potential consequences of climate change for primary production and fish production in large marine ecosystems. Philosophical Transactions of the Royal Society of London, Series B: Biological Sciences, 367, 2979–2989.

Brown, J.H., Gillooly, J.F., Allen, A.P., Savage, V.M. & West, G.B. (2004). Toward a metabolic theory of ecology. Ecology, 85, 1771–1789.

Byrd, R.H., Lu, Peihuang., Nocedal, Jorge. & Zhu, Ciyou. (1995). A Limited Memory Algorithm for Bound Constrained Optimization. SIAM J. Sci. Comput., 16, 1190–1208.

Canales, T.M., Law, R. & Blanchard, J.L. (2016). Shifts in plankton size spectra modulate growth and coexistence of anchovy and sardine in upwelling systems. Canadian Journal of Fisheries and Aquatic Sciences, 73, 611–621.

Casini, M., Hjelm, J., Molinero, J.-C., Lövgren, J., Cardinale, M., Bartolino, V., et al. (2009). Trophic cascades promote threshold-like shifts in pelagic marine ecosystems. Proceedings of the National Academy of Sciences, USA, 106, 197–202.

Casini, M., Käll, F., Hansson, M., Plikshs, M., Baranova, T., Karlsson, O., et al. (2016). Hypoxic areas, density-dependence and food limitation drive the body condition of a heavily exploited marine fish predator. Royal Society Open Science, 3, 160416.

Clarke, A. & Johnston, N.M. (1999). Scaling of metabolic rate with body mass and temperature in teleost fish. Journal of Animal Ecology, 68, 893–905.

Daufresne, M., Lengfellner, K. & Sommer, U. (2009). Global warming benefits the small in aquatic ecosystems. Proceedings of the National Academy of Sciences, USA, 106, 12788–12793.

Dell, A.I., Pawar, S. & Savage, V.M. (2011). Systematic variation in the temperature dependence of physiological and ecological traits. Proceedings of the National Academy of Sciences, 108, 10591–10596.

Dieterich, C., Wang, S., Schimanke, S., Gröger, M., Klein, B., Hordoir, R., et al. (2019). Surface Heat Budget over the North Sea in Climate Change Simulations. Atmosphere, 10, 272.

van Dorst, R.M., Gårdmark, A., Svanbäck, R., Beier, U., Weyhenmeyer, G.A. & Huss, M. (2019). Warmer and browner waters decrease fish biomass production. Global Change Biology, 25, 1395–1408.

Englund, G., Öhlund, G., Hein, C.L. & Diehl, S. (2011). Temperature dependence of the functional response. Ecology Letters, 14, 914–921.

Forster, J., Hirst, A.G. & Atkinson, D. (2012). Warming-induced reductions in body size are greater in aquatic than terrestrial species. PNAS, 109, 19310–19314.

Free, C.M., Thorson, J.T., Pinsky, M.L., Oken, K.L., Wiedenmann, J. & Jensen, O.P. (2019). Impacts of historical warming on marine fisheries production. Science, 363, 979–983.

Galbraith, E.D., Carozza, D.A. & Bianchi, D. (2017). A coupled human-Earth model perspective on long-term trends in the global marine fishery. Nat Commun, 8, 14884.

Gårdmark, A., Casini, M., Huss, M., van Leeuwen, A., Hjelm, J., Persson, L., et al. (2015). Regime shifts in exploited marine food webs: detecting mechanisms underlying alternative stable states using size-structured community dynamics theory. Phil. Trans. R. Soc. B, 370, 20130262.

Gårdmark, A. & Huss, M. (2020). Individual variation and interactions explain food web responses to global warming. Philosophical Transactions of the Royal Society B: Biological Sciences, 375, 20190449.

Gilbert, B., Tunney, T.D., McCann, K.S., DeLong, J.P., Vasseur, D.A., Savage, V.M., et al. (2014). A bioenergetic framework for the temperature dependence of trophic interactions. Ecology Letters, 17, 902–914.

Gillooly, J.F., Brown, J.H., West, G.B., Savage, V.M. & Charnov, E.L. (2001). Effects of size and temperature on metabolic rate. Science, 2248–2251.

Gröger, M., Arneborg, L., Dieterich, C., Höglund, A. & Meier, H.E.M. (2019). Summer hydrographic changes in the Baltic Sea, Kattegat and Skagerrak projected in an ensemble of climate scenarios downscaled with a coupled regional ocean–sea ice– atmosphere model. Clim Dyn, 53, 5945–5966.

Guiet, J., Aumont, O., Poggiale, J.-C. & Maury, O. (2016). Effects of lower trophic level biomass and water temperature on fish communities: A modelling study. Progress in Oceanography, 146, 22–37.

Handeland, S.O., Imsland, A.K. & Stefansson, S.O. (2008). The effect of temperature and fish size on growth, feed intake, food conversion efficiency and stomach evacuation rate of Atlantic salmon post-smolts. Aquaculture, 283, 36–42.

Hartvig, M., Andersen, K.H. & Beyer, J.E. (2011). Food web framework for size-structured populations. Journal of Theoretical Biology, 272, 113–122.

Heneghan, R.F., Galbraith, E., Blanchard, J.L., Harrison, C., Barrier, N., Bulman, C., et al. (2021). Disentangling diverse responses to climate change among global marine ecosystem models. Progress in Oceanography, 198, 102659.

Heneghan, R.F., Hatton, I.A. & Galbraith, E.D. (2019). Climate change impacts on marine ecosystems through the lens of the size spectrum. Emerging Topics in Life Sciences, 3, 233–243.

Hilborn, R. & Walters, C.J. (1992). Quantitative Fisheries Stock Assessment: Choice, Dynamics and Uncertainty. Springer, Norwell MA, USA.

Huss, M., Lindmark, M., Jacobson, P., van Dorst, R.M. & Gårdmark, A. (2019). Experimental evidence of gradual size-dependent shifts in body size and growth of fish in response to warming. Glob Change Biol, 25, 2285–2295.

ICES. (2013). Report of the Baltic Fisheries Assessment Working Group (WGBFAS) (No. ICES CM 2013/ACOM:10.). 10-17 April 2013 ICES Headquarters, Copenhagen.

ICES. (2015). Report of the Baltic Fisheries Assessment Working Group (WGBFAS) (No. ICES CM 2015/ACOM:10). 14-21 April 2015 ICES Headquarters, Copenhagen.

ICES. (2021). Report of the Baltic Fisheries Assessment Working Group (WGBFAS) (No. 3:53).

Ikpewe, I.E., Baudron, A.R., Ponchon, A. & Fernandes, P.G. (2020). Bigger juveniles and smaller adults: Changes in fish size correlate with warming seas. Journal of Applied Ecology, 58, 847–856.

Jacobsen, N.S., Burgess, M.G. & Andersen, K.H. (2017). Efficiency of fisheries is increasing at the ecosystem level. Fish and Fisheries, 18, 199–211.

Jennings, S. & Collingridge, K. (2015). Predicting Consumer Biomass, Size-Structure, Production, Catch Potential, Responses to Fishing and Associated Uncertainties in the World’s Marine Ecosystems. PLOS ONE, 10, e0133794.

Jorgensen, C., Enberg, K., Dunlop, E.S., Arlinghaus, R., Boukal, D.S., Brander, K., et al. (2007). Ecology: managing evolving fish stocks. Science, 318, 1247–1248.

Kwiatkowski, L., Torres, O., Bopp, L., Aumont, O., Chamberlain, M., Christian, J.R., et al. (2020). Twenty-first century ocean warming, acidification, deoxygenation, and upper-ocean nutrient and primary production decline from CMIP6 model projections. Biogeosciences, 17, 3439–3470.

Lefort, S., Aumont, O., Bopp, L., Arsouze, T., Gehlen, M. & Maury, O. (2015). Spatial and body-size dependent response of marine pelagic communities to projected global climate change. Global Change Biology, 21, 154–164.

Lindmark, M., Ohlberger, J. & Gårdmark, A. (2022). Optimum growth temperature declines with body size within fish species. Global Change Biology, 28, 2259–2271.

Lorenzen, K. (2016). Toward a new paradigm for growth modeling in fisheries stock assessments: Embracing plasticity and its consequences. Fisheries Research, Growth: theory, estimation, and application in fishery stock assessment models, 180, 4–22.

Lotze, H.K., Tittensor, D.P., Bryndum-Buchholz, A., Eddy, T.D., Cheung, W.W.L., Galbraith, E.D., et al. (2019). Global ensemble projections reveal trophic amplification of ocean biomass declines with climate change. Proceedings of the National Academy of Sciences, 116, 12907–12912.

Mackenzie, B.R., Gislason, H., Möllmann, C. & Köster, F.W. (2007). Impact of 21st century climate change on the Baltic Sea fish community and fisheries. Global Change Biology, 13, 1348–1367.

McCauley, D.J., Pinsky, M.L., Palumbi, S.R., Estes, J.A., Joyce, F.H. & Warner, R.R. (2015). Marine defaunation: Animal loss in the global ocean. Science, 347.

McCoy, M.W. & Gillooly, J.F. (2008). Predicting natural mortality rates of plants and animals. Ecology Letters, 11, 710–716.

Mion, M., Haase, S., Hemmer-Hansen, J., Hilvarsson, A., Hüssy, K., Krüger-Johnsen, M., et al. (2021). Multidecadal changes in fish growth rates estimated from tagging data: A case study from the Eastern Baltic cod (Gadus morhua, Gadidae). Fish and Fisheries, 22, 413–427.

Mion, M., Thorsen, A., Vitale, F., Dierking, J., Herrmann, J.P., Huwer, B., et al. (2018). Effect of fish length and nutritional condition on the fecundity of distressed Atlantic cod Gadus morhua from the Baltic Sea: POTENTIAL FECUNDITY OF BALTIC G. MORHUA. Journal of Fish Biology, 92, 1016–1034.

Möllmann, C., Diekmann, R., Müller-Karulis, B., Kornilovs, G., Plikshs, M. & Axe, P. (2009). Reorganization of a large marine ecosystem due to atmospheric and anthropogenic pressure: a discontinuous regime shift in the Central Baltic Sea. Global Change Biology, 15, 1377–1393.

Morán, X.A.G., López-Urrutia, Á., Calvo-Díaz, A. & Li, W.K.W. (2010). Increasing importance of small phytoplankton in a warmer ocean. Global Change Biology, 16, 1137–1144.

Morita, K., Fukuwaka, M., Tanimata, N. & Yamamura, O. (2010). Size-dependent thermal preferences in a pelagic fish. Oikos, 119, 1265–1272.

Munch, S.B. & Salinas, S. (2009). Latitudinal variation in lifespan within species is explained by the metabolic theory of ecology. Proceedings of the National Academy of Sciences, 106, 13860–13864.

Neubauer, P. & Andersen, K.H. (2019). Thermal performance of fish is explained by an interplay between physiology, behaviour and ecology. Conserv Physiol, 7.

Neuenfeldt, S., Bartolino, V., Orio, A., Andersen, K.H., Andersen, N.G., Niiranen, S., et al. (2020). Feeding and growth of Atlantic cod (Gadus morhua L.) in the eastern Baltic Sea under environmental change. ICES Journal of Marine Science, 77, 624–632.

Neuheimer, A.B. & Grønkjaer, P. (2012). Climate effects on size-at-age: growth in warming waters compensates for earlier maturity in an exploited marine fish. Global Change Biology, 18, 1812–1822.

Neuheimer, A.B., Thresher, R.E., Lyle, J.M. & Semmens, J.M. (2011). Tolerance limit for fish growth exceeded by warming waters. Nature Climate Change, 1, 110–113.

Ohlberger, J. (2013). Climate warming and ectotherm body size – from individual physiology to community ecology. Functional Ecology, 27, 991–1001.

Ohlberger, J., Edeline, E., Vollestad, L.A., Stenseth, N.C. & Claessen, D. (2011). Temperature-driven regime shifts in the dynamics of size-structured populations. The American Naturalist, 177, 211–223.

Pauly, D. (1980). On the interrelationships between natural mortality, growth parameters, and mean environmental temperature in 175 fish stocks. ICES Journal of Marine Science, 39, 175–192.

Pinsky, M.L., Worm, B., Fogarty, M.J., Sarmiento, J.L. & Levin, S.A. (2013). Marine Taxa Track Local Climate Velocities. Science, 341, 1239–1242.

Pontavice, H. du, Gascuel, D., Reygondeau, G., Maureaud, A. & Cheung, W.W.L. (2019). Climate change undermines the global functioning of marine food webs. Global Change Biology.

R Core Team. (2020). R: A Language and Environment for Statistical Computing. R Foundation for Statistical Computing. Vienna, Austria.

Rall, B.C., Brose, U., Hartvig, M., Kalinkat, G., Schwarzmuller, F., Vucic-Pestic, O., et al. (2012). Universal temperature and body-mass scaling of feeding rates. Philosophical Transactions of the Royal Society of London, Series B: Biological Sciences, 367, 2923–2934.

Reum, J.C.P., Blanchard, J.L., Holsman, K.K., Aydin, K. & Punt, A.E. (2019). Species-specific ontogenetic diet shifts attenuate trophic cascades and lengthen food chains in exploited ecosystems. Oikos, 128, 1051–1064.

Righton, D.A., Andersen, K.Haste., Neat, F., Thorsteinsson, V., Steingrund, P., Svedäng, H., et al. (2010). Thermal niche of Atlantic cod Gadus morhua: limits, tolerance and optima. Marine Ecology Progress Series, 420, 1–13.

van Rijn, I., Buba, Y., DeLong, J., Kiflawi, M. & Belmaker, J. (2017). Large but uneven reduction in fish size across species in relation to changing sea temperatures. Global Change Biology, 23, 3667–3674.

Rohatgi, A. (2012). WebPlotDigitalizer: HTML5 based online tool to extract numerical data from plot images. Version 4.1. [WWW document] URL https://automeris.io/WebPlotDigitizer (accessed on January 2019).

Savage, V.M., Gillooly, J.F., Brown, J.H., West, G.B. & Charnov, E.L. (2004). Effects of body size and temperature on population growth. The American Naturalist, 163, 429–441.

Scott, F., Blanchard, J. & Andersen, K. (2019). mizer: Multi-Species sIZE Spectrum Modelling in R. R.

Scott, F., Blanchard, J.L. & Andersen, K.H. (2014). mizer: An R package for multispecies, trait-based and community size spectrum ecological modelling. Methods in Ecology and Evolution, 5, 1121–1125.

Scott, F., Blanchard, J.L. & Andersen, K.Haste. (2018). Multispecies, trait and community size spectrum ecological modelling in R (mizer), 1–87.

Sheridan, J.A. & Bickford, D. (2011). Shrinking body size as an ecological response to climate change. Nature Climate Change, 1, 401–406.

Steinacher, M., Joos, F., Frolicher, T.L., Bopp, L., Cadule, P., Cocco, V., et al. (2010). Projected 21st century decrease in marine productivity: a multi-model analysis. Biogeosciences, 7.

Svedäng, H. & Hornborg, S. (2014). Selective fishing induces density-dependent growth. Nature Communications, 5, 4152.

Szuwalski, C.S., Burgess, M.G., Costello, C. & Gaines, S.D. (2017). High fishery catches through trophic cascades in China. Proceedings of the National Academy of Sciences, 114, 717–721.

Thorson, J.T., Monnahan, C.C. & Cope, J.M. (2015). The potential impact of time-variation in vital rates on fisheries management targets for marine fishes. Fisheries Research, 169, 8–17.

Thorson, J.T., Munch, S.B., Cope, J.M. & Gao, J. (2017). Predicting life history parameters for all fishes worldwide. Ecological Applications, 27, 2262–2276.

Thresher, R.E., Koslow, J.A., Morison, A.K. & Smith, D.C. (2007). Depth-mediated reversal of the effects of climate change on long-term growth rates of exploited marine fish. Proceedings of the National Academy of Sciences, USA, 104, 7461–7465.

Tittensor, D.P., Novaglio, C., Harrison, C.S., Heneghan, R.F., Barrier, N., Bianchi, D., et al. (2021). Next-generation ensemble projections reveal higher climate risks for marine ecosystems. Nat. Clim. Chang., 11, 973–981.

Tu, C.-Y., Chen, K.-T. & Hsieh, C. (2018). Fishing and temperature effects on the size structure of exploited fish stocks. Sci Rep, 8, 7132.

Ursin, E. (1967). A Mathematical Model of Some Aspects of Fish Growth, Respiration, and Mortality. Journal of the Fisheries Research Board of Canada, 24, 2355–2453.

Ursin, E. (1973). On the prey size preferences of cod and dab. Meddelelser fra Danmarks Fiskeri-og Havundersgelser, 7:8598.

Woodworth-Jefcoats, P.A., Blanchard, J.L. & Drazen, J.C. (2019). Relative Impacts of Simultaneous Stressors on a Pelagic Marine Ecosystem. Frontiers in Marine Science, 6.

Woodworth-Jefcoats, P.A., Polovina, J.J., Dunne, J.P. & Blanchard, J.L. (2013). Ecosystem size structure response to 21st century climate projection: large fish abundance decreases in the central North Pacific and increases in the California Current. Global Change Biology, 19, 724–733.

Woodworth-Jefcoats, P.A., Polovina, J.J., Howell, E.A. & Blanchard, J.L. (2015). Two takes on the ecosystem impacts of climate change and fishing: Comparing a size-based and a species-based ecosystem model in the central North Pacific. Progress in Oceanography, 138, 533–545.

Yvon-Durocher, G., Montoya, J.M., Trimmer, M. & Woodward, G. (2011). Warming alters the size spectrum and shifts the distribution of biomass in freshwater ecosystems. Global Change Biology, 17, 1681–1694.

