## Supporting information for "Temperature impacts on fish physiology and resource abundance lead to faster growth but smaller fish sizes and yields under warming"

### Model parameterization

#### Species-specific parameters

Below follows a description for how we acquired default parameters which are valid at the reference temperature, $T_{ref}$, where the temperature scaling equals 1 for all rates (see main text for which rates are assumed temperature dependent and how). In *'mizer'*, multi-species size spectrum models (MSSMs), species-specific parameters are required. Parameters that have no defaults and must be provided are: asymptotic size ($W$), size at maturation ($w_{mat}$), preferred predator-prey mass ratio (PPMR) ($\beta$), the standard deviation of PPMR ($\sigma$) and the maximum recruitment in the Beverton-Holt stock recruitment function ($R_{max}$) (Scott *et al.* 2019).

For cod, we initially acquired the parameters $\beta$ and $\sigma$ by calculating the mean and the standard deviation for the distribution of mean individual-level log PPMR ($\beta$ is then exponentiated whereas $\sigma$ is defined as the standard deviation of the distribution of log PPMRs), using stomach data (downloaded on 2019.03.20) hosted by the International Council for the Exploration of the Sea (ICES), in the FISH STOMACH database (ICES 2010). These data can be downloaded at http://ecosystemdata.ices.dk/stomachdata/download.aspx. We used only samples with fully intact prey (no digestion) among the observations from the area we consider (ICES subdivisions SD 25-29, 32, see Fig. S2) and from the time window we use for calibrating the model (1992-2002). When prey weight was not available, we used the relationship $weight = 0.01 \times length^3$, where length is in cm, to approximate weight in g. However, since stomach content data is not a measure of preference per se, as it is only shows what has been eaten (which also depends on prey availability), these parameters should also be evaluated after model fitting. For sprat and herring, we assumed $\beta$ and $\sigma$ to be 1000 and 1.3, respectively, which reflects that forage fish typically feed on prey that are small in relation to their body size throughout ontogeny (Aydin *et al.* 2002; Reum *et al.* 2019).

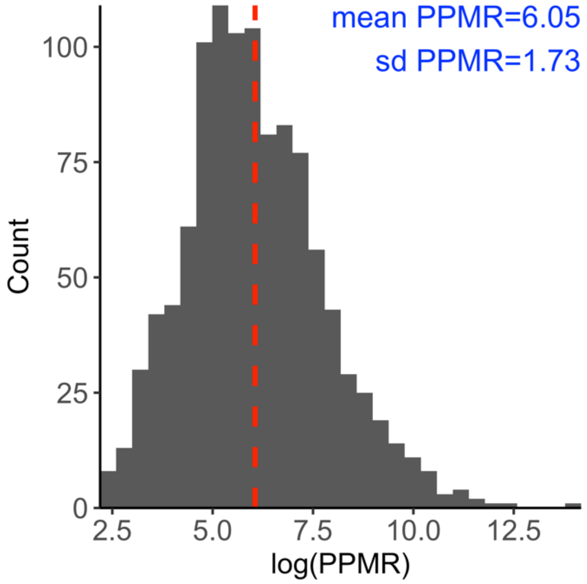

*Figure S1. Distribution of individual level predator-prey mass ratio (*log *PPMR) for cod. Mean* exp*(PPMR) corresponds to $\beta$ and sd to $\sigma$. The dashed red vertical line corresponds to the mean* log *PPMR.*

From von Bertalanffy (VBGE) growth parameters and theory linking growth to feeding parameters (Hartvig *et al.* 2011), the remaining species-specific parameters can be estimated. The allometric constant in the maximum food intake rate ($h_i$) is defined as $h_i = \frac{3k_{vb}}{\alpha f_0} W^{1/3}$, where $k_{vb}$ is the Brody growth coefficient of the VBGE, $\alpha$ is the assimilation efficiency, $f_0$ is the initial feeding level of small individuals and $W$ is asymptotic mass (see main text) (Andersen *et al.* 2009; Scott *et al.* 2019) (but see calibration protocol). When $h_i$ is known, the allometric constant in the search rate function, $\gamma_i$, can be calculated (see equation 9 in main text). The allometric constant of standard metabolism, $k_{met,i}$ is by default $0.12h_i$ (but see calibration protocol).

We estimated VBGE parameters $k_{vb}$ and $W$ for cod, herring and sprat in the Baltic Sea using data from the Baltic International Trawl Survey (BITS), maintained by ICES. They are publicly available at the DATRAS database (http://www.ices.dk/marine-data/data-portals/Pages/DATRAS.aspx). We downloaded the full data for the years 1992-2002 (the calibration time period) on 2018.11.24, and applied further processing in R using the '*tidyverse*' packages (Wickham *et al.* 2019). We followed the approach in Ogle (2013) and fit the VBGE using the form $L_a = L_\infty(1 - e^{-k_{vb}(a-t_0)})$ when estimating parameters. Here $L_a$ is the expected length at age $a$, $L_\infty$ is the average asymptotic length, $k_{vb}$ is the Brody growth rate coefficient and $t_0$ is a modeling artifact that represents the time or age when the average length would have been zero. Parameters were estimated using non-linear least squares regression ('*nls*' package in R), and the packages '*FSA*' (Ogle 2018) and '*FSAdata*' (Ogle 2017). We fit VBGE parameters using length-at-age data and convert between length and weight using the equation $w = aL^b$, where parameters $a$ and $b$ are estimated in this study from a subset of the BITS data where both weight and length information is available for the same individual. Length-at-maturation was taken from the literature and was converted to weight-at-maturation using length-weight relationships estimated in this study from BITS (Table S1). All models were fitted using R (R Core Team 2020), and for all statistical models we verified through visual inspection that assumptions about error homoscedasticity and normality were met.

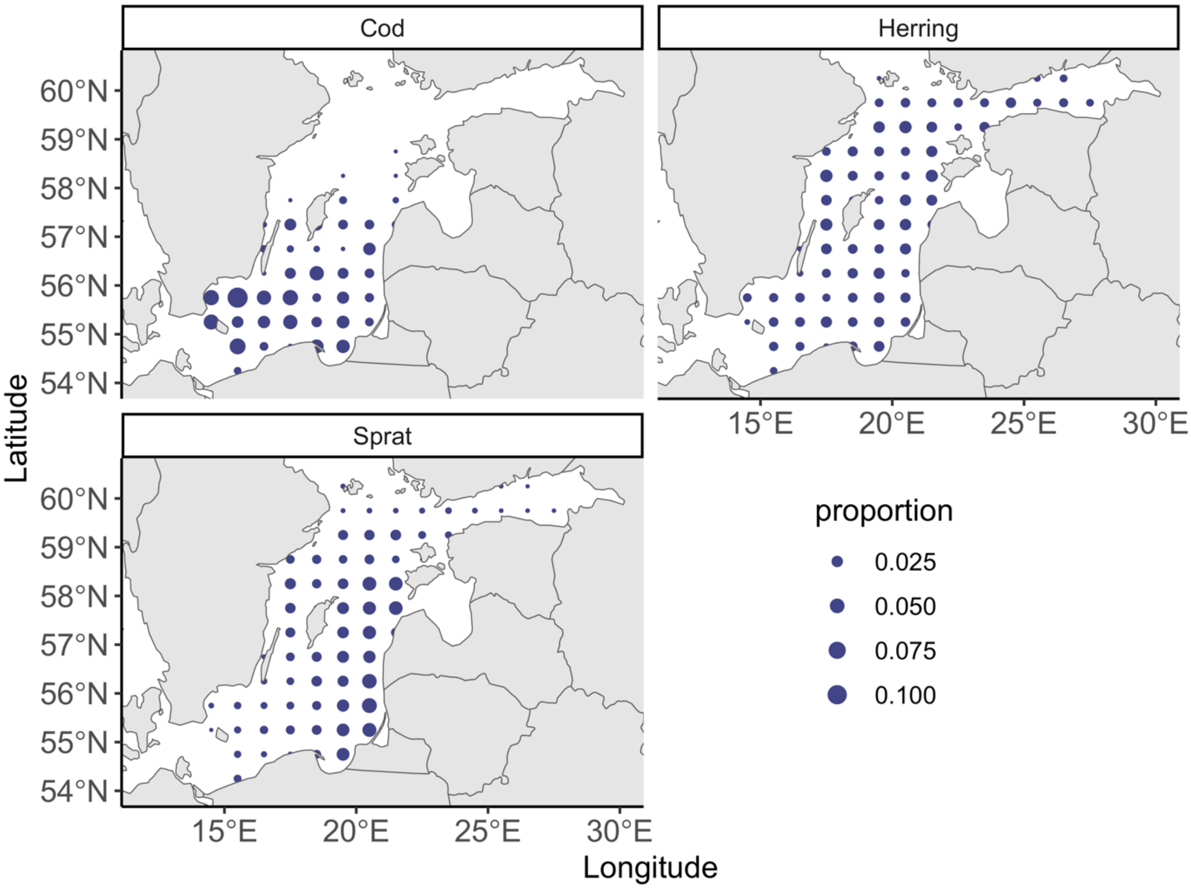

*Figure S2. Map of the study area (ICES sub-divisions 25-29+32). Point size is proportional to abundance per ICES rectangle during the calibration time period (1992-2002), using data from the Baltic International Trawl Survey (BITS) for cod, and the Baltic International Acoustic Survey (BIAS) for sprat and herring.*

*Table S1. Species-specific parameters used in model. Source: 1 = estimated in this study, 2 = (Reum et al. 2019), 3 = (Hartvig et al. 2011), 4 = generalized values based on (Casini et al. 2004), 5 = calculated internally. Parameters in bold are tuned in the calibration process, parenthesis show default values pre-calibration.*

| Symbol | Description | Unit | Cod | Herring | Sprat | Source |
|---|---|---|---|---|---|---|
| $\boldsymbol{w_{mat}}$ | Maturation size* | g | 267 | 17 | 4 | 1 |
| $\boldsymbol{W}$ | Asymptotic weight | g | 44903 | 125 | 21 | 1 |
| $\boldsymbol{k_{vb}}$ | Brody growth coefficient | $\mathrm{yr}^{-1}$ | 0.07 | 0.165 | 0.287 | 1 |
| $\boldsymbol{t_0}$ | Age at length 0 | - | -0.94 | -3.74 | -2.97 | 1 |
| $\boldsymbol{\beta}$ | Preferred predator prey mass ratio | - | 426 | 1000 | 1000 | 1,2 |
| $\boldsymbol{\sigma}$ | Width of prey size preference | - | 1.7 | 1.3 | 1.3 | 3 |
| $\boldsymbol{h}$ | Constant for max. food intake | $\mathrm{g}^{1-n}\ \mathrm{yr}^{-1}$ | **27 (20.7)** | **8.9 (6.9)** | **8.6 (6.6)** | 5 |
| $\boldsymbol{\gamma}$ | Constant for volumetric search rate | $\mathrm{g}^{-q}\mathrm{m}^2\ \mathrm{yr}^{-1}$ | **0.6 (0.38)** | **0.18 (0.11)** | **0.17 (0.1)** | 5 |
| $\boldsymbol{k_{met}}$ | Constant for metabolic rate (note this term is assumed to represent all metabolic costs, e.g., standard and for activity) | $\mathrm{g}^{1-p}\ \mathrm{yr}^{-1}$ | 2.49 | 0.83 | 0.22 | 5 |
| $\boldsymbol{\epsilon, erepro}$ | Reproductive efficiency | - | **5e-05 (0.01)** | **8e-04 (0.01)** | **1e-03 (0.01)** | 1 |
| $\boldsymbol{a}$ | Constant in length-weight relationship | - | 0.0078 | .0042 | .0041 | 1 |
| $\boldsymbol{b}$ | Exponent in length-weight relationship | - | 3.07 | 3.14 | 3.15 | 1 |
| $\boldsymbol{R_{max}}$ | **Maximum recruitment in the Beverton-Holt stock recruitment function**** | $\mathrm{g/m}^2$ | **0.003 (0.0826)** | **0.1 (11.1)** | **1.16 (7.3)** | **-** |
| $\boldsymbol{\theta_{i,ben}}$ | Benthos availability | - | 0.5 | 0.5 | 0 | 4 |
| $\boldsymbol{\theta_{i,pla}}$ | Plankton availability | - | 0.5 | 0.5 | 1 | 4 |

*Length at maturation for cod and sprat are taken from van Leeuwen *et al*. (2013) (30 cm and 9 cm, respectively) and for herring we used Huss *et al*. (2012) (14 cm), based on (Vainikka *et al*. 2009a, b). **See calibration protocol for unit and volume scaling

*Table S2. General parameters. Source: 1 = (Scott et al. 2019), 2 = (Hartvig et al. 2011), 3 = (Blanchard et al. 2014), 4 = this study, 5 = Audzijonyte et al., unpublished.. Parameters in bold are tuned in the calibration process, parenthesis show default values pre-calibration.*

| Symbol | Description | Value | Unit | Source |
|---|---|---|---|---|
| $\boldsymbol{\alpha}$ | Assimilation efficiency | 0.6 | - | 1,2 |
| $\boldsymbol{w_0}$ | Egg weight | 0.001 | g | 1,2 |
| $\boldsymbol{f_0}$ | Initial feeding level | 0.6 | - | 1,2 |
| $\boldsymbol{n}$ | Exponent of max. consumption | $2/3$ | - | 1,2 |
| $\boldsymbol{m}$ | Exponent to $\frac{w}{w_{mat,i}}$ in smooth maturation function | 5 | - | 5, 4 |
| $\boldsymbol{q}$ | Exponent of search volume | 0.8 | - | 3 |
| $\boldsymbol{p}$ | Exponent of metabolic rate | 0.7 | - | 3 |
| $\boldsymbol{\mu_0}$ | Pre-factor for background mortality | 0.6 | $\mathrm{yr}^{-1}$ | 1 |
| $\boldsymbol{\lambda}$ | Exponent of background resource spectra | $-2-q+n$ | - | 2 |
| $\boldsymbol{r_0}$ | Regeneration rate of background resource spectra | 4 | $\mathrm{g}^{1-\mathrm{n}}\mathrm{yr}^{-1}$ | 4 |
| $\boldsymbol{\kappa}$ | **Carrying capacity of background resource spectra (benthos, plankton)** | **9 (11)** | $\mathbf{g^{\lambda-1}m^2}$ | **4** |
| $\boldsymbol{w_{cut,P}}$ | Cut-off size of plankton size spectrum | 1 | g | 4 |
| $\boldsymbol{w_{cut,B}}$ | Cut-off size of benthic size spectrum | 20 | g | 4 |

#### Temperature dependence

*Table S3 Parameters of distributions describing activation energies of temperature-dependent rates in the size spectrum model. See Figure S11 for 200 random draws from these normal distributions that were used as input in projections.*

| **Symbol** | **Rate** | **Distribution (mean, s.d)** | **Source** |
|---|---|---|---|
| $\boldsymbol{A_{met}}$ | Metabolism | $Normal(0.62, 0.025)$ | (Lindmark *et al.* 2022) |
| $\boldsymbol{A_{mor}}$ | Background mortality | $Normal(0.62, 0.025)$ | (Pauly 1980; Brown *et al.* 2004; Lindmark *et al.* 2022) |
| $\boldsymbol{A_{int}}$ | Maximum consumption; Search volume* | $Normal(0.69, 0.078)$ | (Lindmark *et al.* 2022) |
| $\boldsymbol{A_{gro}}$ | Background resources regeneration rate | $Normal(0.73, 0.1)$ | Experimental data from Savage *et al.* (2004) for algae, phyto- and zooplankton. |
| $\boldsymbol{A_{car}}$ | Background resource carrying capacity | $Normal(-0.8, 0.1)$ | Assumed to be $-\boldsymbol{A_{gro}}$, based on metabolic arguments and constant resources (Savage *et al.* 2004; Gilbert *et al.* 2014; Bernhardt *et al.* 2018) |

* For simplicity we assume maximum consumption to scale the same way with temperature as search volume, and the parameter estimates come from maximum consumption experiments.

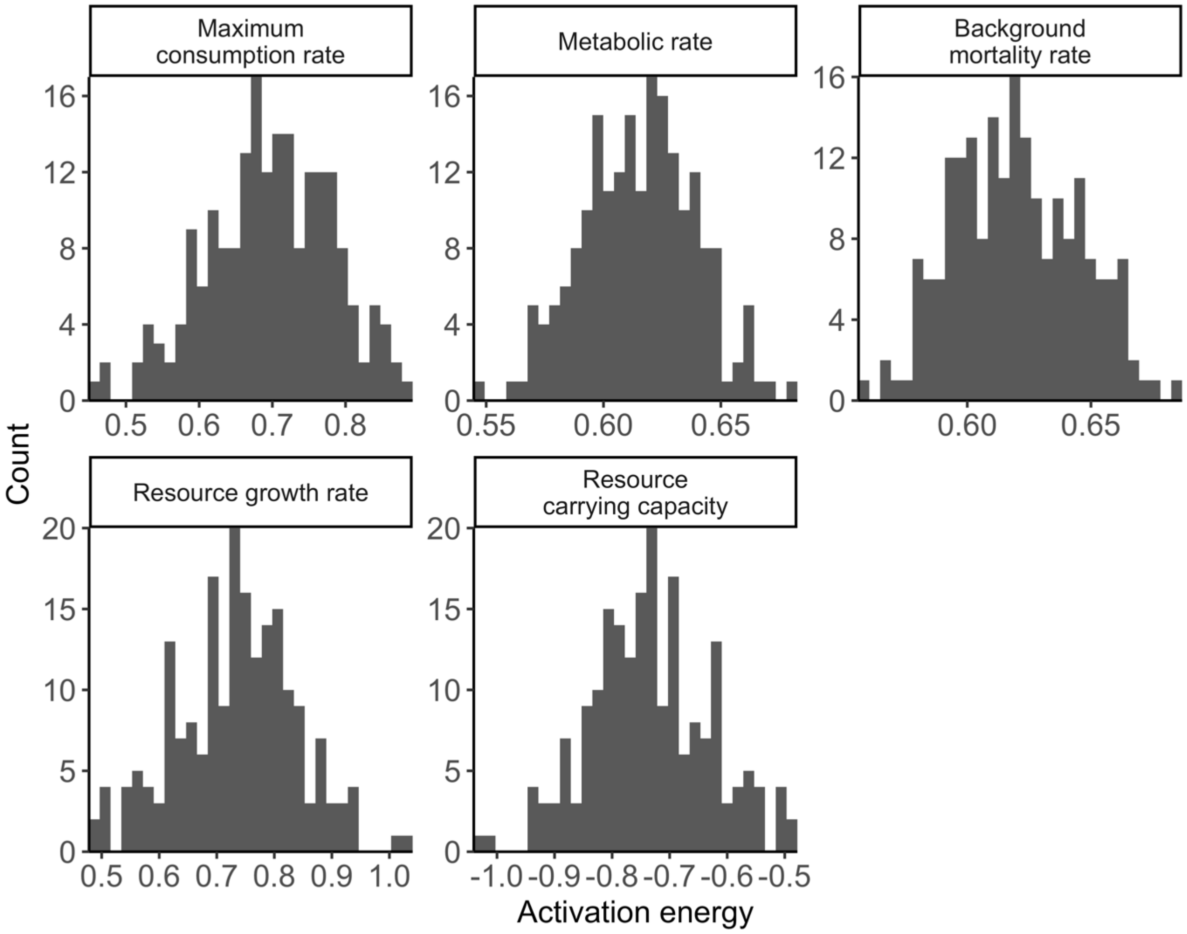

*Figure S3. 200 random samples from the distributions describing the activation energies of the rates resource carrying capacity and resource growth rates (assumed same for benthic and pelagic background resources), metabolic rate, background mortality rates and maximum consumption rates. See Table S3. Combinations of these parameters were used in model projections to understand how uncertainty in these parameters affect individual- and population level metrics.*

### Model calibration and validation

#### Calibration protocol

We calibrated to the model using mean SSB and F between 1992-2002 from stock assessments, which were acquired from ICES Working Group Reports (ICES 2013, 2015), and are in unit $10^6$ kg/area. This area corresponds to ICES subdivisions 25-29+32, which equals approximately $2.49\times10^{11}$ $m^2$ (estimated using ICES shapefiles) (ICES 2021). We first defined the size spectrum model in units $g/m^2$ and then used the conversion factor 249 to express biomasses in unit 1000 tonnes/area. Below follows a step-by-step description of how the model was calibrated to the assembled Baltic Sea data after being initially parameterized.

1. Determine a starting value for plankton and benthos $\kappa$. The aim is to find coexistence of the three fish species and for their spawning stock biomasses (SSBs) to be within an order of magnitude of SSBs from stock assessment. Prioritize coexistence over fits of SSBs, as these are calibrated later. As starting values for $R_{max,i}$, which ensures coexistence, we used the same ratio between $R_{max,i}$ and $\kappa$ as calibrated in (Blanchard et al. 2014), expressed in unit $m^2$ (by dividing with $10^{11}$, which corresponds roughly to the area of the North Sea). This procedure resulted in a starting value for $\kappa_B$ and $\kappa_P$ of 11, in unit $g^{\lambda-1}m^2$.
2. Evaluate body growth rates against empirical data.
    a. If modelled body growth rates are low, check the species-specific feeding levels $f_i(w)$. The feeding level describes the level of satiation, with 0 being unfed and 1 completely satiated. For reference, a feeding level of 0.2 is the minimum to cover basic metabolic costs (by default) and thus does not allow for body growth, whereas a constant feeding level of 0.6 fits a von Bertalanffy curve (Andersen *et al.* 2009; Scott *et al.* 2019). If low feeding level seem likely as a cause for poor growth in the model, check the search rate ($\gamma$) or available prey. If not, check the net energy acquisition.
        i. Net energy acquisition is largely determined by maximum consumption and metabolic losses. Earlier explorations suggest modelled growth rates can be lower than observed (Scott *et al.* 2019) with default $k_{met}$ and $h$ (constants in allometric metabolism and maximum consumption

rates, respectively), so these may need to be modified based on empirical data or other theoretically derived relationships.

ii. Go back to step 1. With altered $k_{met}$ and/or $h$, $\kappa$ needed for coexistence and fit to SSB may change. Therefore, a $\kappa$ can be chosen that puts modelled SSBs closer to SSBs from stock assessment, while still allowing for coexistence and maintaining modelled growth rates in line with observed.

3. If $\kappa$ allows for coexistence and results in SSBs within an order of magnitude to SSBs obtained from stock assessments and body growth rates are realistic, tune $R_{max}$ to match SSBs from the size spectrum model closer to that from the stock assessment. This is done by finding the vector of $R_{max}$-values that minimizes the residual sum of squares between the two SSBs.
4. Verify that SSBs and growth rates are still close to empirical data, after optimizing $R_{max}$. Then evaluate the ratio of egg production before and after density dependence is added to the Beverton-Holt type stock-recruit curve (RDI/RDD).
   a. If RDI/RDD is high ($\approx 100$) (Jacobsen *et al.* 2017), and RDD is close to $R_{max}$, consider lowering $erepro$, because the high RDI/RDD indicates that species are resistant to fishing (flat yield~fishing mortality curves).
      i. Go back to Step 2 and proceed forward from there.
5. Verify that SSBs, growth rates and RDI/RDD are realistic. Then assess emergent diets.
   a. If predatory interactions do not match independent stomach data, re-evaluate the predation kernel. We judged this for cod based on (Niiranen *et al.* 2019), who show that benthos (*Saduria entomon*) are found in stomachs of small cod whereas sprat and herring start to appear in the size classes of 20-29 cm. We assume sprat and herring feed mostly on background spectra with only limited piscivory and cannibalism.
6. Project forward and backwards with time-varying fishing mortality from stock assessment output and temperature data. Compare qualitative trends with stock assessment data. Since the stock assessment data for the three fish species are outputs from different models and assumptions, this serves more as a useful comparison rather than validation of the model.

#### Results from calibration procedure

The following figures show model output from the final set of parameters found through the calibration procedure described above (Table S1-S2). In addition to the bold parameters in Tables S1-S2 ($\epsilon$ and $R_{max}$), we also updated the constant in the maximum consumption rate ($h$) with a factor of 1.3 as body growth was otherwise poor despite a normal feeding level (~0.6). The level of density dependence imposed by the stock-recruit function (see Eq. 14-15) was evaluated by assessing the ratio of the physiological recruitment, $R_{phy,i}$, to the recruitment $R_i$ (RDI/RDD in the *mizer* package) (Jacobsen *et al*. 2017). If this ratio is 1, there is no additional density dependence from the stock-recruitment function because the species are at the linear part of the stock-recruit relationship. After calibrating the model, we acquired $\frac{R_{phy,i}}{R_i}$ ratios of 554, 79 and 28 for cod, herring and sprat, respectively. High values (>100) suggest recruitment is largely controlled by the external parameter $R_{max}$ (Jacobsen *et al.* 2017), which in our case can be said for cod, and to a lesser extent, sprat and herring.

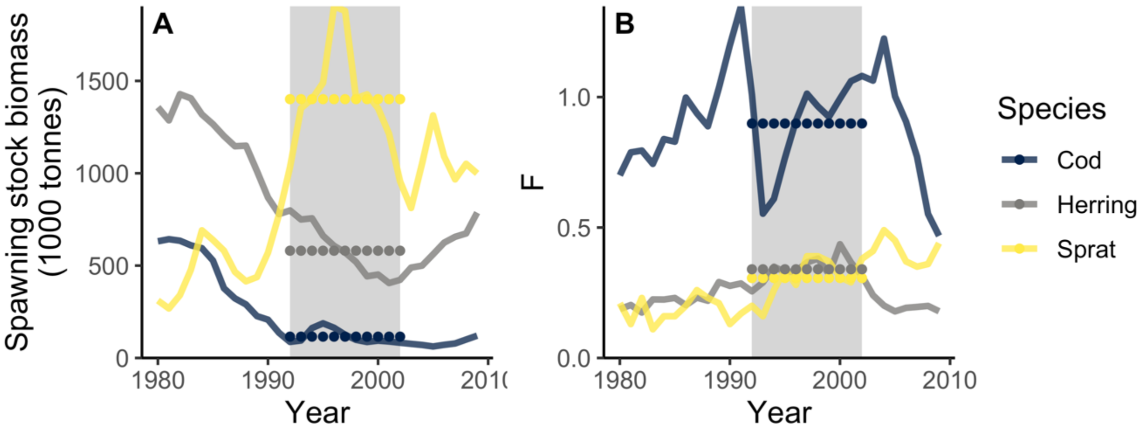

*Figure S4. Time-series of spawning stock biomass (SSB) (A) and fishing mortality (B) from stock assessment model estimates (ICES 2013, 2015). Grey background shows calibration period, dotted horizontal lines show the mean of each time series in the calibration time period.*

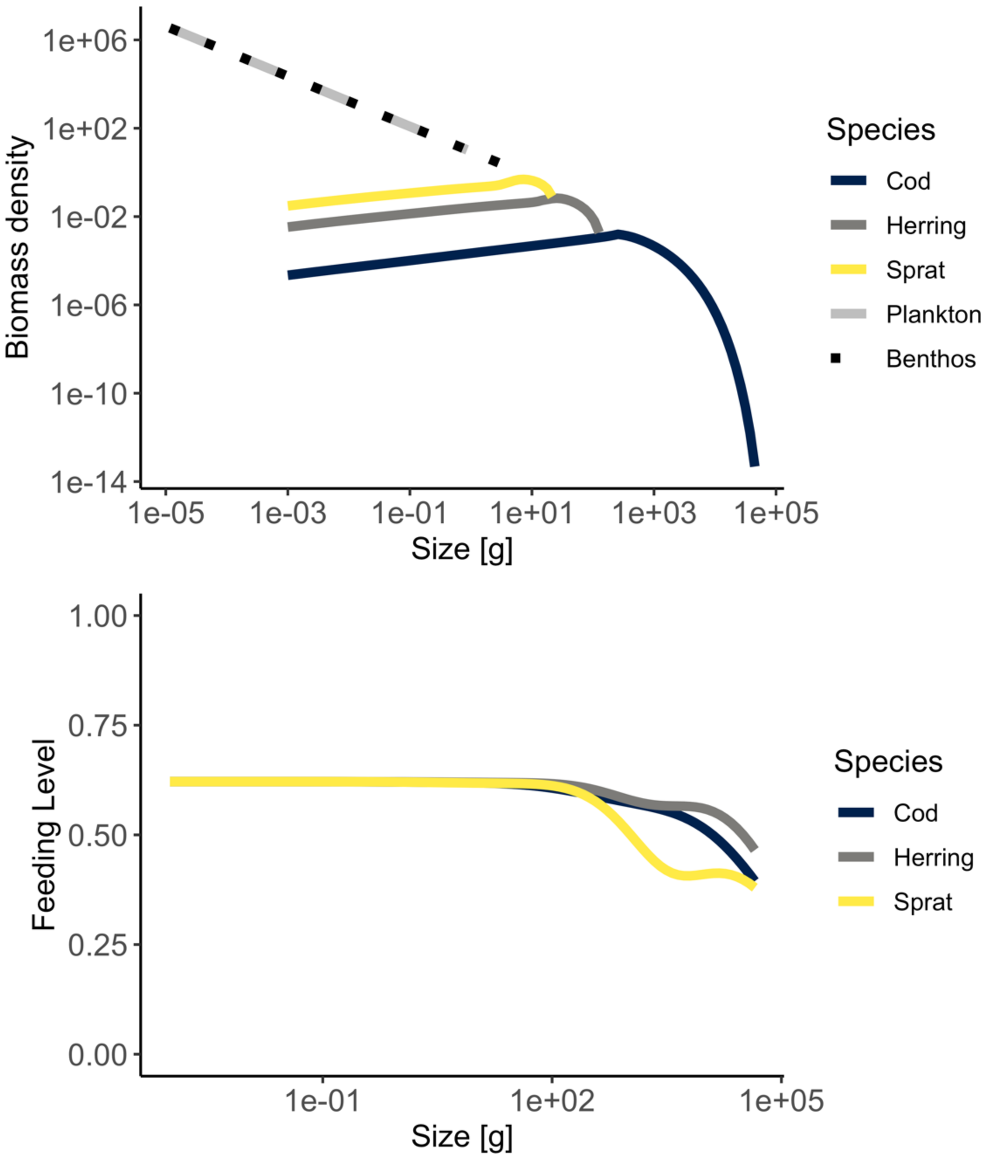

*Figure S5. Biomass spectra of species and background resources (top) and feeding levels as a function of body size (g) from the size spectrum model simulated to steady state using the calibrated model parameters (Table S1-S2).*

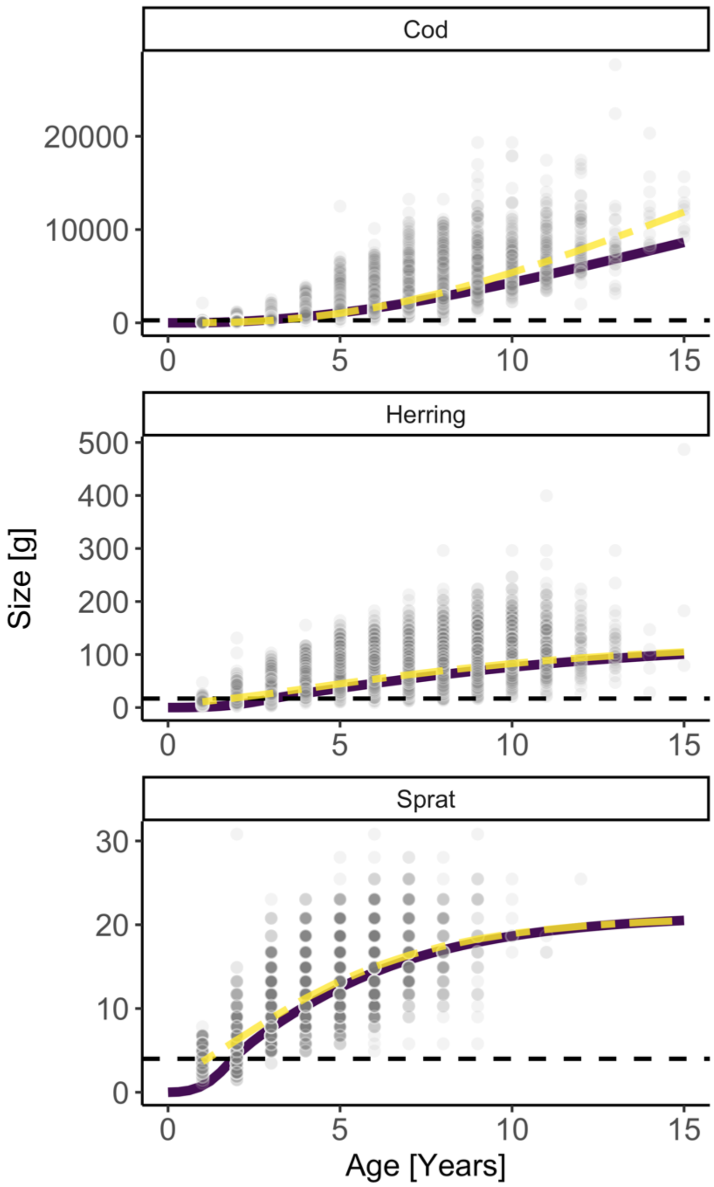

*Figure S6. Size-at-age (g) from size spectrum model (solid purple line) run to steady state using the calibrated model parameters, growth curves from the von Bertalanffy growth equation (two-dashed yellow line) fitted to length-at-age data from the Baltic International Trawl Survey (semitransparent dark grey points) and then converted to mass using the length-weight relationship estimated in this study.*

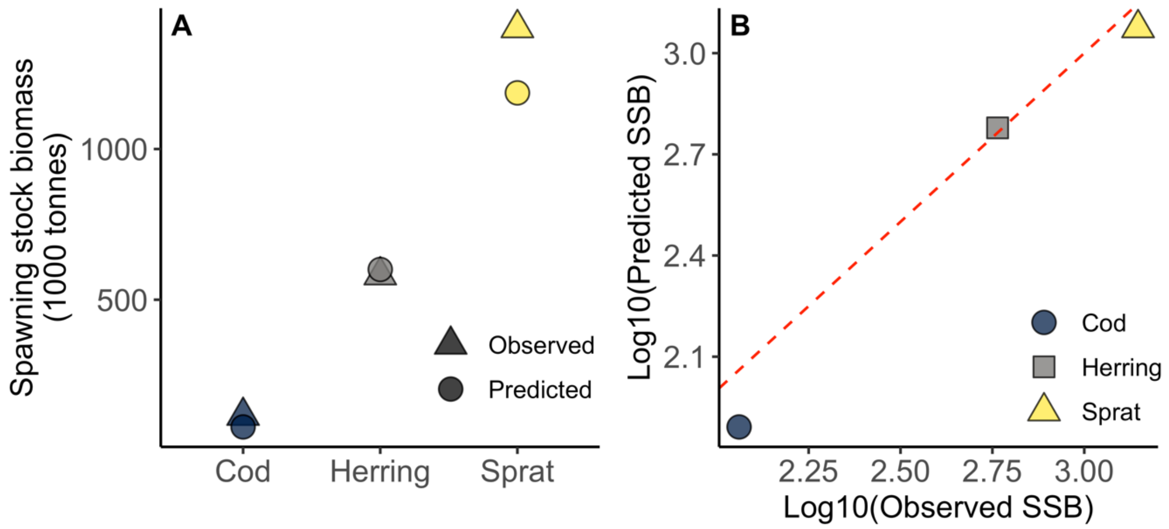

*Figure S7. Predicted (size-spectrum model) vs 'observed' (stock assessment) spawning stock biomass (SSB) per species. A) Mean predicted SSB (final 20 time steps of projection) for each species and observed SSB in the calibration period. B) Same data on log10 scale plotted against each other with a 1:1 line (red dashed).*

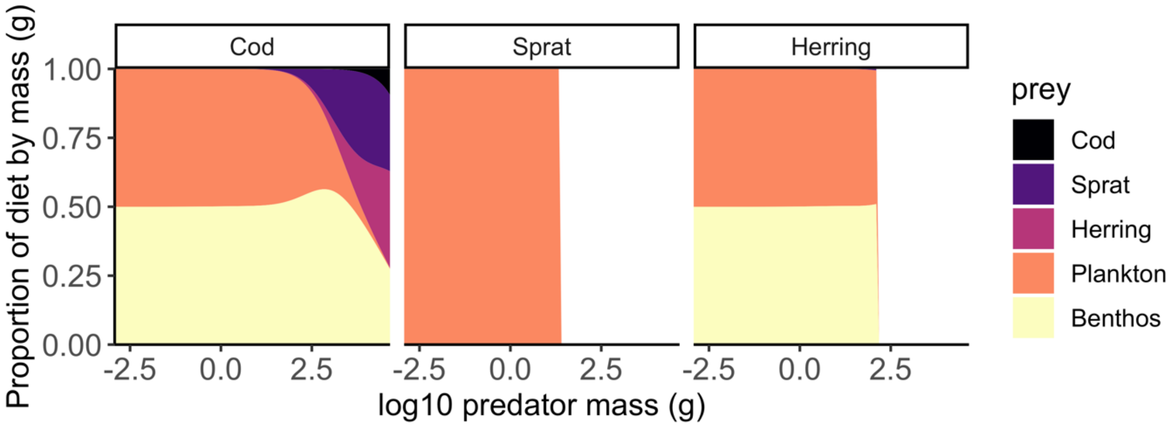

*Figure S8. Proportion of diet by mass as a function of predator body mass at steady state. Cod becomes piscivorous when approximately 25cm (Niiranen* et al. *2019). This roughly corresponds to a mass of about 150g, which is slightly smaller than what emerges from the model (~300 g).*

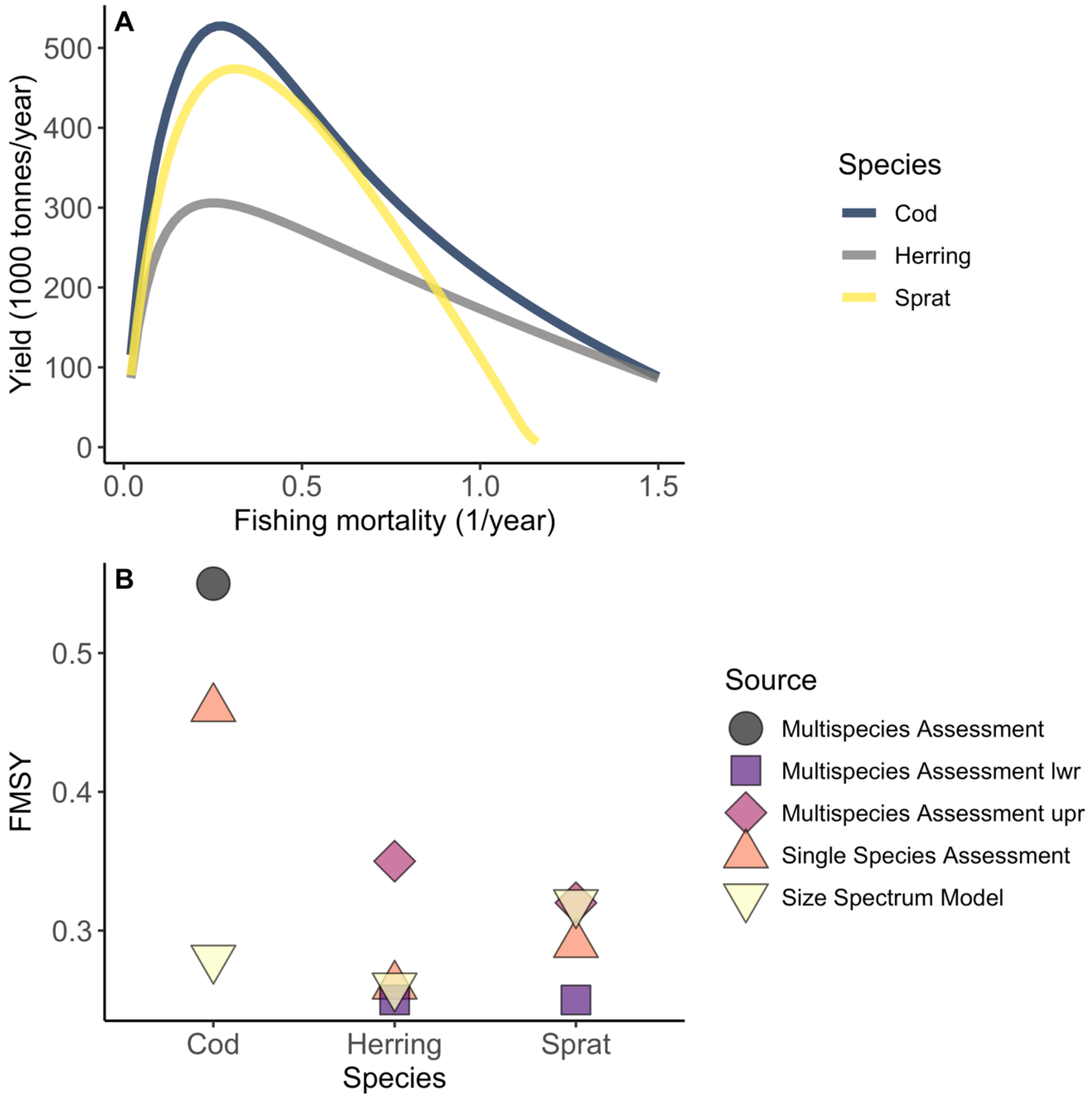

*Figure S9. (A) Yield in the size spectrum model (estimated by keeping each species at their mean assessment $F_{MSY}$) at the reference temperature ($T_{ref}$) and (B) corresponding $F_{MSY}$ (fishing mortality resulting in the highest long-term yield) for each species in the size spectrum model compared to those obtained in multi- and single-species stock assessment models (ICES 2013, 2015), using default parameters.*

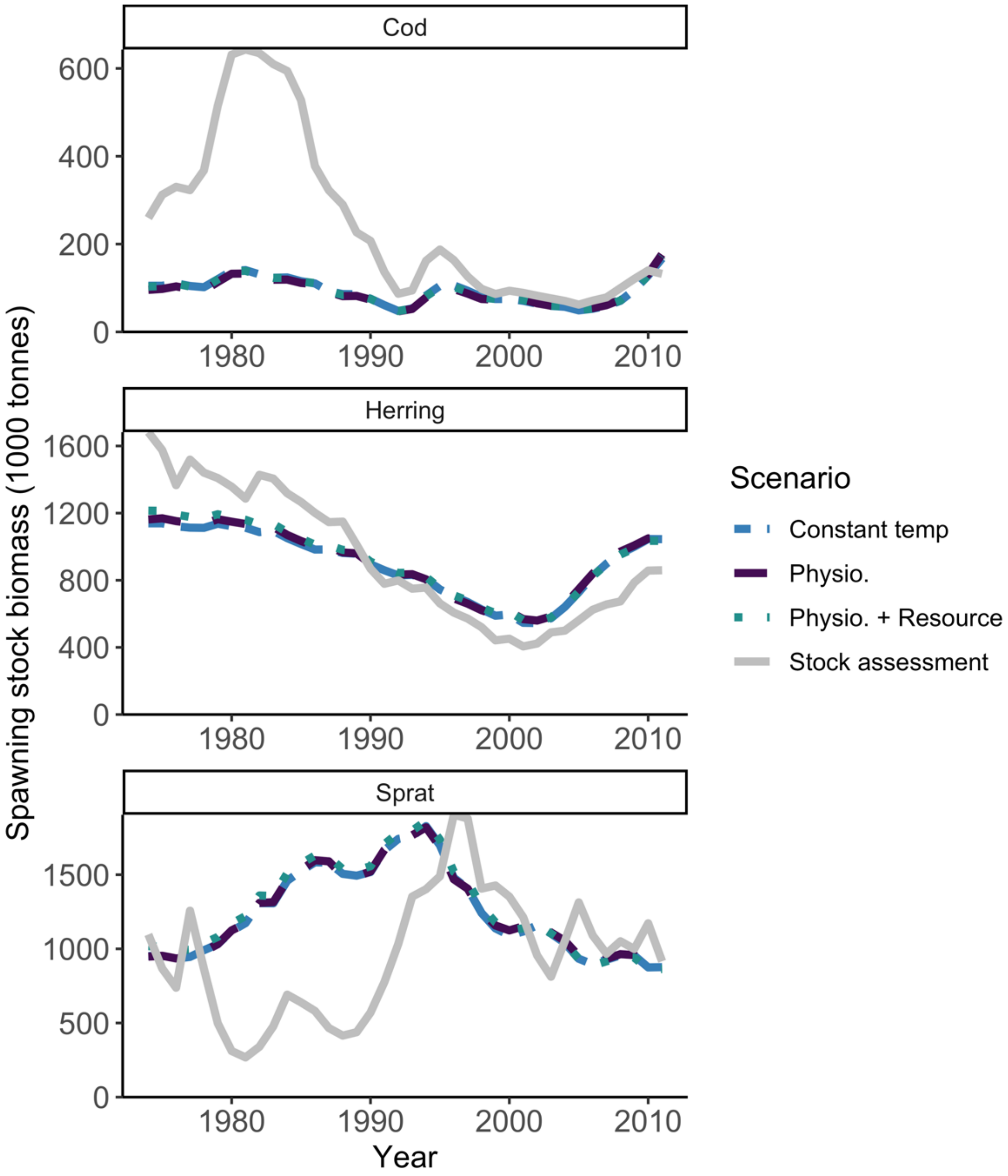

*Figure S10. Temporal development of spawning stock biomass (SSB). Temperature varies in the time series according to the RCA4-NEMO model using the RCP 8.5 scenario. The projections start in 1874 (not shown) to allow a 100-year burn-in period. This is sufficient to reach steady state before historical fishing efforts are introduced (in 1974-2012). F-values by species in the burn-in period are equal to the values in 1974 (first year of effort-time series). The solid grey lines are SSB from stock assessments, the two-dashed blue lines are SSBs from the size-spectrum model projections assuming constant temperatures (equivalent to no temperature effects). The purple dashed line is the model where only physiological processes are temperature dependent, the dotted teal line is where physiological processes and resources are temperature dependent. Mean activation energies are used (Table S3).*

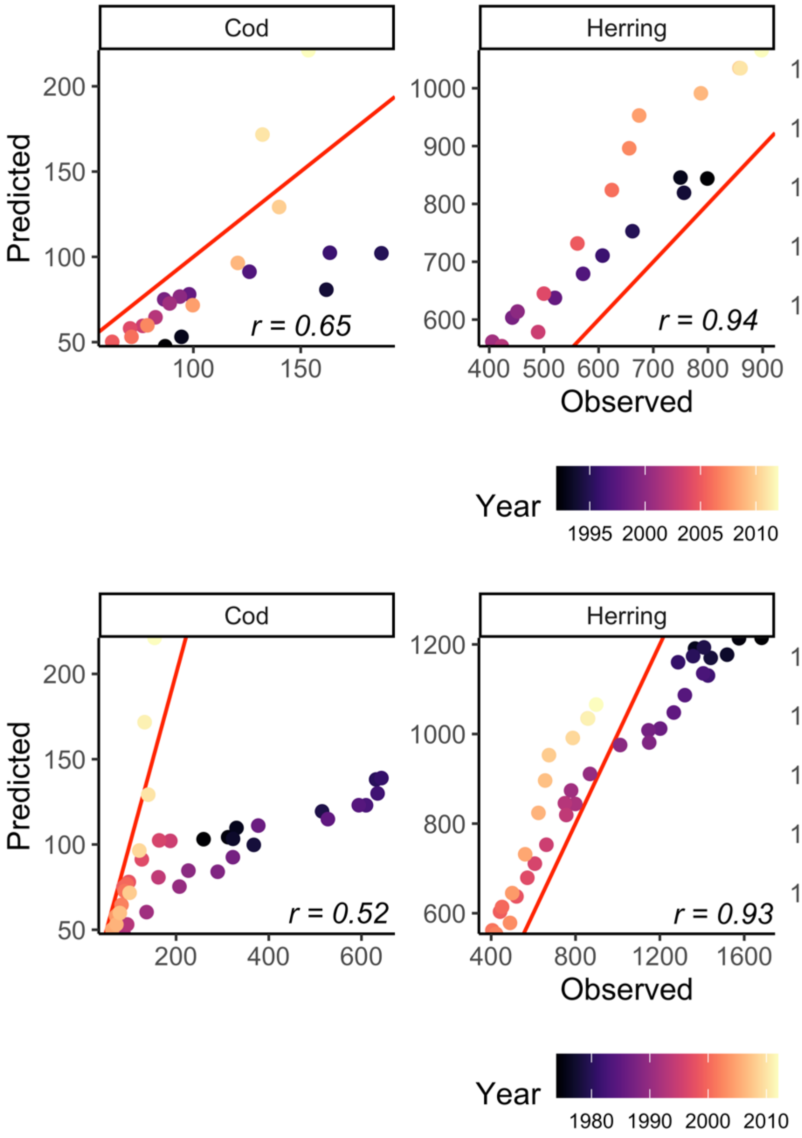

*Figure S11. Correlation between SSB from size-spectrum model ("predicted") and stock assessment models ("observed") between 1992-2012, i.e., from the start of the calibration to the last year (top row) and 1974-2012, i.e., from the start of the time series (bottom row). The Pearson correlation coefficients are indicated in the bottom right corner. Colors depict year.*

### Analysis

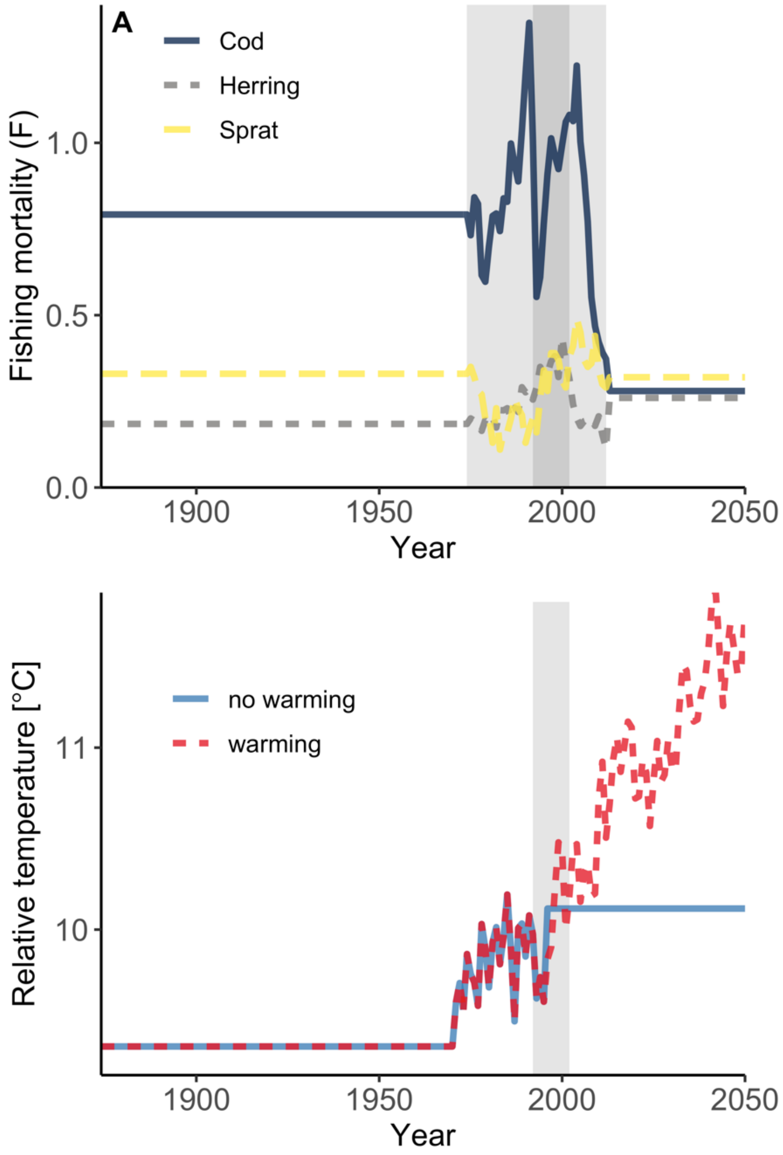

*Figure S12. Time series of (A) fishing mortality by species and (B) relative temperature. For fishing mortality (A) the light grey area corresponds to years where mortality values are estimated from stock assessments. The dark grey area is the model calibration time period. We treated the 100 years prior to the start of the F-time series as burn-in and kept the F constant and equal to the first F time series (in year 1974). The future projections correspond to the* $F_{MSY}$ *estimated in the size spectrum model (estimated by keeping each species at their mean assessment* $F_{MSY}$*). The relative temperature is scaled by adding a constant of 10.11562 to the relative change in sea surface temperature from the regional coupled model system RCA4-NEMO using the RCP 8.5. This constant makes the mean temperature in the calibration period the same as the reference temperature,* $T_{ref}$ *(i.e., no temperature effects). Years prior to the temperature model are assigned the first value in the temperature record. In all time-varying*

*temperature projections, the initial temperature is the same. The no-warming scenario gets a constant temperature equal to the mean temperature in the calibration time-period in 1997 (blue solid line, mid-year in calibration time window) while the warming scenario continuous along the temperature projection from the regional coupled model system RCA4-NEMO using the RCP 8.5 scenario (red dashed line).*

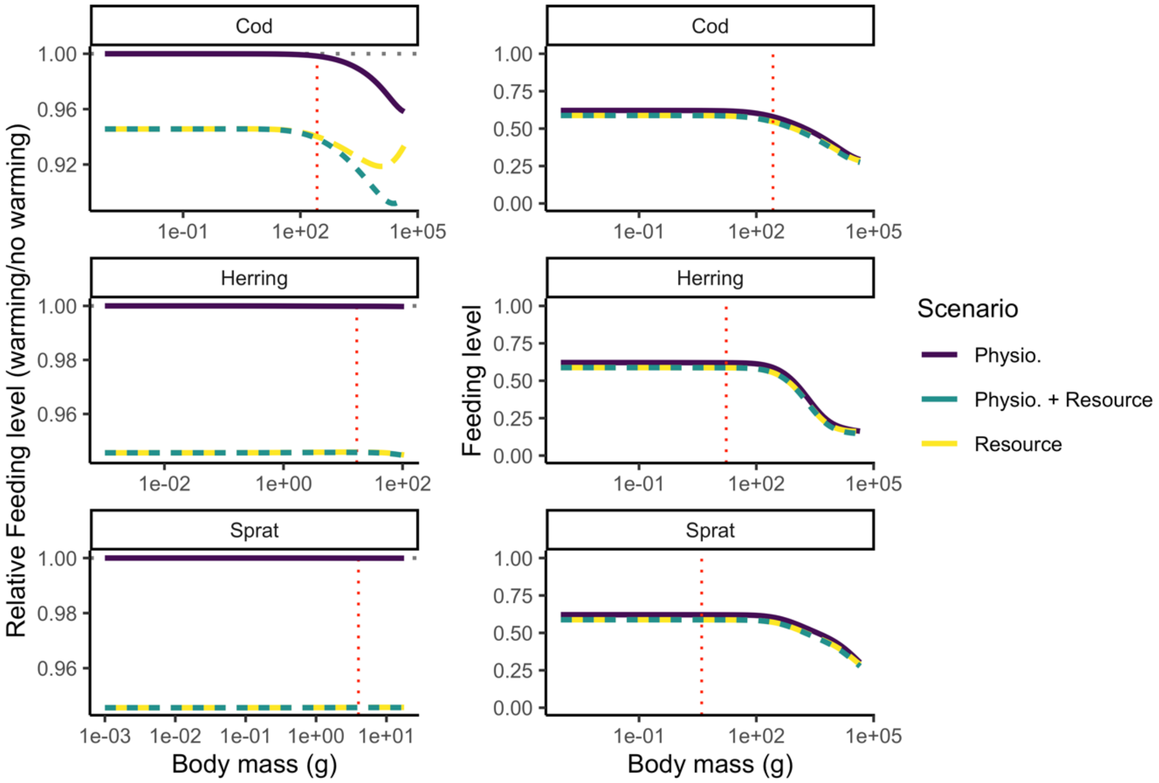

*Figure S13. Feeding levels from model projections to 2050 assuming warming according to RCP 8.5 while keeping fishing mortality at average* $F_{MSY}$ *levels (as estimated at* $T_{ref}$*). Left column shows feeding levels relative to a non-warming simulation (i.e., dashed horizontal grey line) and right column shows absolute feeding level. Activation energies are the means of their respective distributions (no uncertainty).*

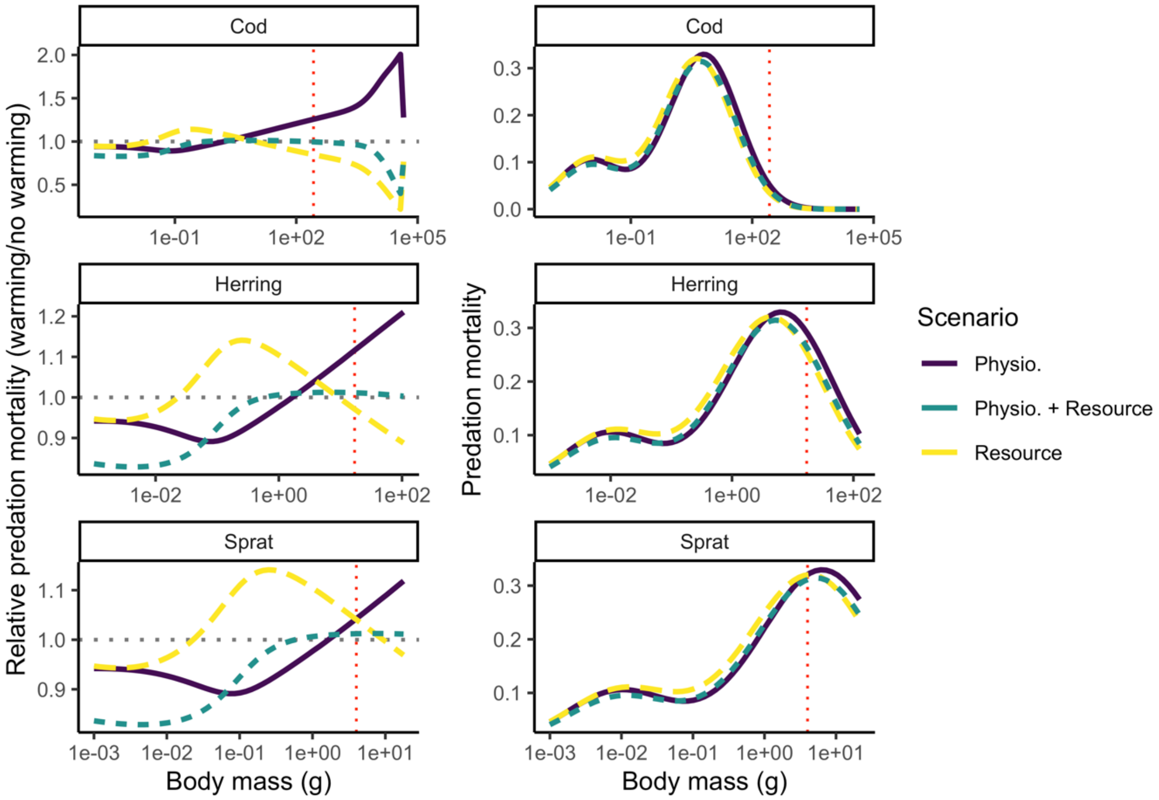

*Figure S14. Feeding levels from model projections to 2050 assuming warming according to RCP 8.5 while keeping fishing mortality at average $F_{MSY}$ levels (as estimated at $T_{ref}$). Left column shows predation mortality levels relative to a non-warming simulation (i.e., dashed horizontal grey line) and right column shows absolute predation mortality. Activation energies are the means of their respective distributions (no uncertainty).*

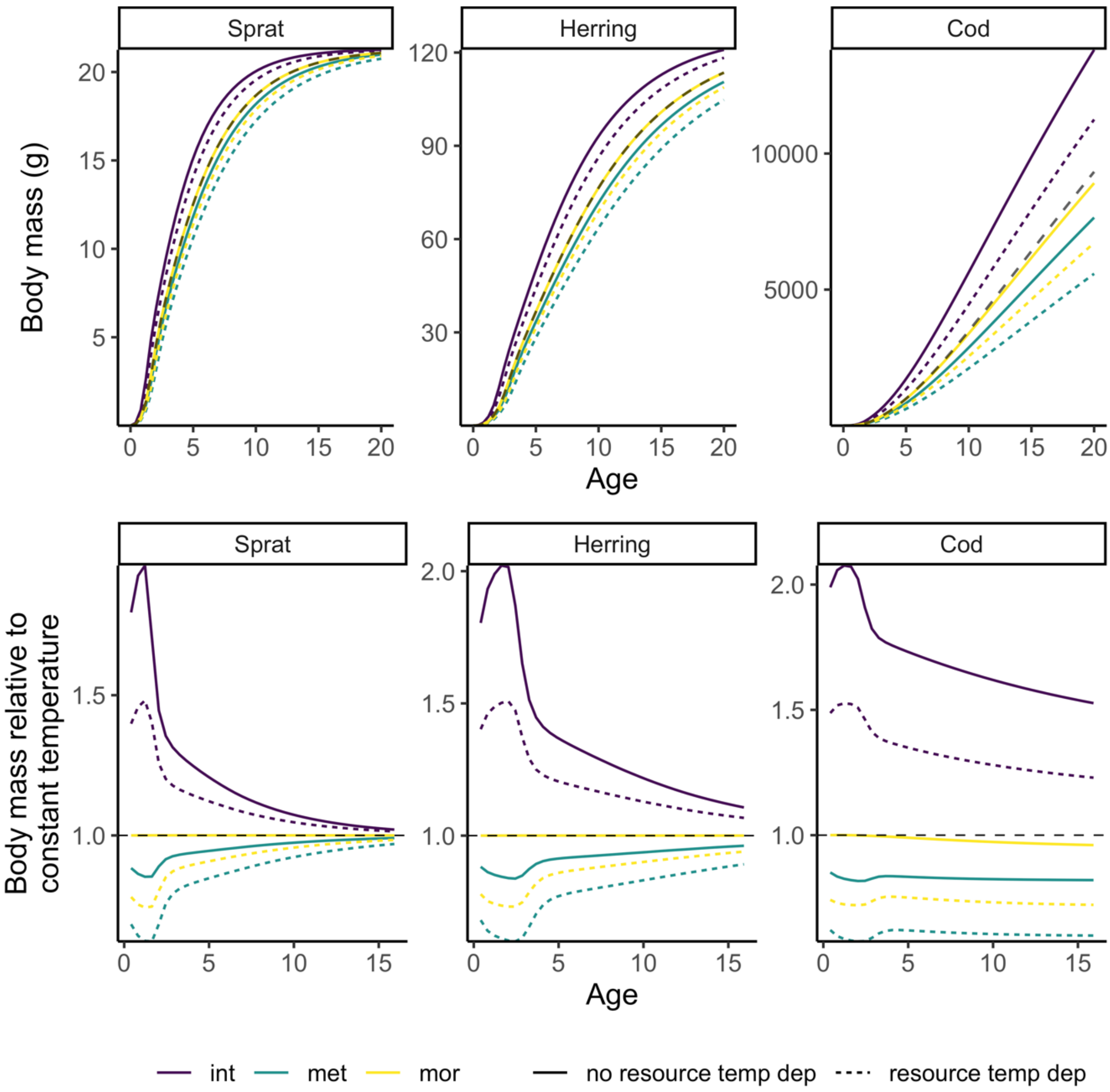

*Figure S15. The effect of including temperature dependence for one physiological process for growth with temperatures $T_{ref} + 2°C$. Here we include temperature-dependence for one physiological process at the time (purple=only intake rates, green=only metabolic rates, yellow=only mortality rates), with and without temperature dependence on resources (dashed and solid lines, respectively). Top panel depicts size-at-age and the bottom panel depicts size-at-age relative to size-at-age from a projection at $T_{ref}$ non-temperature dependent model.*

### References

Andersen, K.H., Farnsworth, K.D., Pedersen, M., Gislason, H. & Beyer, J.E. (2009). How community ecology links natural mortality, growth, and production of fish populations, 7.

Aydin, K.Y., Lapko, V.V., Radchenko, V.I. & Livingston, P.A. (2002). *A Comparison of the Eastern and Western Bering Sea Shelf/Slope Ecosystems Through the Use of Mass Balance Food Web Models. - US Dept of Commerce, NOAA Techical. Memo. NMFS-AFSC-130, 78*.

Bernhardt, J.R., Sunday, J.M. & O'Connor, M.I. (2018). Metabolic Theory and the Temperature-Size Rule Explain the Temperature Dependence of Population Carrying Capacity. *The American Naturalist*, 192, 687–697.

Blanchard, J.L., Andersen, K.H., Scott, F., Hintzen, N.T., Piet, G. & Jennings, S. (2014). Evaluating targets and trade-offs among fisheries and conservation objectives using a multispecies size spectrum model. *Journal of Applied Ecology*, 51, 612–622.

Blanchard, J.L., Jennings, S., Holmes, R., Harle, J., Merino, G., Allen, J.I., *et al.* (2012). Potential consequences of climate change for primary production and fish production in large marine ecosystems. *Philosophical Transactions of the Royal Society of London, Series B: Biological Sciences*, 367, 2979–2989.

Brown, J.H., Gillooly, J.F., Allen, A.P., Savage, V.M. & West, G.B. (2004). Toward a metabolic theory of ecology. *Ecology*, 85, 1771–1789.

Casini, M., Cardinale, M. & Arrhenius, F. (2004). Feeding preferences of herring () and sprat () in the southern Baltic Sea. *ICES Journal of Marine Science*, 61, 1267–1277.

Gilbert, B., Tunney, T.D., McCann, K.S., DeLong, J.P., Vasseur, D.A., Savage, V.M., *et al.* (2014). A bioenergetic framework for the temperature dependence of trophic interactions. *Ecology Letters*, 17, 902–914.

Hartvig, M., Andersen, K.H. & Beyer, J.E. (2011). Food web framework for size-structured populations. *Journal of Theoretical Biology*, 272, 113–122.

Huss, M., Gårdmark, A., van Leeuwen, A. & de Roos, A.M. (2012). Size- and food-dependent growth drives patterns of competitive dominance along productivity gradients. *Ecology*, 93, 847–857.

ICES. (2010). Stomach Dataset 2010, ICES, Copenhagen.

ICES. (2013). *Report of the Baltic Fisheries Assessment Working Group (WGBFAS)* ( No. ICES CM 2013/ACOM:10.). 10-17 April 2013 ICES Headquarters, Copenhagen.

ICES. (2015). *Report of the Baltic Fisheries Assessment Working Group (WGBFAS)* ( No. ICES CM 2015/ACOM:10). 14-21 April 2015 ICES Headquarters, Copenhagen.

ICES. (2021). *ICES Spatial Facility, ICES, Copenhagen*.

Jacobsen, N.S., Burgess, M.G. & Andersen, K.H. (2017). Efficiency of fisheries is increasing at the ecosystem level. *Fish and Fisheries*, 18, 199–211.

van Leeuwen, A., Huss, M., Gårdmark, A., Casini, M., Vitale, F., Hjelm, J., *et al.* (2013). Predators with multiple ontogenetic niche shifts have limited potential for population growth and top-down control of their prey. *American Naturalist*, 182, 53–66.

Lindmark, M., Ohlberger, J. & Gårdmark, A. (2022). Optimum growth temperature declines with body size within fish species. *Global Change Biology*, 28, 2259–2271.

Möllmann, C., Diekmann, R., Müller-Karulis, B., Kornilovs, G., Plikshs, M. & Axe, P. (2009). Reorganization of a large marine ecosystem due to atmospheric and anthropogenic pressure: a discontinuous regime shift in the Central Baltic Sea. *Global Change Biology*, 15, 1377–1393.

Niiranen, S., Orio, A., Bartolino, V., Bergström, U., Kallasvuo, M., Neuenfeldt, S., *et al.* (2019). Predator-prey body size relationships of cod in a low-diversity marine system. *Mar. Ecol. Prog. Ser.*, 627, 201–206.

Ogle, D.H. (2013). fishR Vignette - Von Bertalanffy Growth Models.

Ogle, D.H. (2017). *FSAdata: Fisheries Stock Analysis, Datasets*.

Ogle, D.H. (2018). *FSA: Fisheries Stock Analysis. R package*.

Pauly, D. (1980). On the interrelationships between natural mortality, growth parameters, and mean environmental temperature in 175 fish stocks. *ICES Journal of Marine Science*, 39, 175–192.

R Core Team. (2020). *R: A Language and Environment for Statistical Computing. R Foundation for Statistical Computing*. Vienna, Austria.

Reum, J.C.P., Blanchard, J.L., Holsman, K.K., Aydin, K. & Punt, A.E. (2019). Species-specific ontogenetic diet shifts attenuate trophic cascades and lengthen food chains in exploited ecosystems. *Oikos*, 128, 1051–1064.

Savage, V.M., Gillooly, J.F., Brown, J.H., West, G.B. & Charnov, E.L. (2004). Effects of Body Size and Temperature on Population Growth. *The American Naturalist*, 163, 429–441.

Scott, F., Blanchard, J. & Andersen, K. (2019). *mizer: Multi-Species sIZE Spectrum Modelling in R*. R. .

Vainikka, A., G\a ardmark, A., Bland, B. & Hjelm, J. (2009a). Two-and three-dimensional maturation reaction norms for the eastern Baltic cod, Gadus morhua. *ICES Journal of Marine Science: Journal du Conseil*, 66, 248–257.

Vainikka, A., Mollet, F., Casini, M. & Gårdmark, A. (2009b). Spatial variation in growth, condition and maturation reaction norms of the Baltic herring Clupea harengus membras. *Marine Ecology Progress Series*, 383, 285–294.

Wickham, H., Averick, M., Bryan, J., Chang, W., D'Agostino McGowan, L., François, R., *et al.* (2019). Welcome to the tidyverse. *Journal of Open Source Software*, 1686.
